# HP1α depletion and TGFβ activation exert antagonistic effects on 3D genome organization

**DOI:** 10.64898/2026.02.12.705282

**Authors:** Fabiana Patalano, Oda Hovet, Roberto Rossini, Maxim Nekrasov, Yasmin Dijkwel, Bhumika Azad, Tatiana Soboleva, Rein Aasland, David Tremethick, Jonas Paulsen

## Abstract

The three-dimensional (3D) organization of the human genome plays a critical role in regulating gene expression and is frequently disrupted in cancer. However, how key factors like Heterochromatin Protein 1 alpha (HP1α) and Transforming Growth Factor beta (TGFβ) remodel this architecture to drive tumorigenesis remains poorly understood. We investigated the effects of HP1α knockdown and TGFβ treatment on higher-order chromatin structure and gene expression in human mammary epithelial cells.

Our findings reveal that HP1α depletion and TGFβ stimulation exert distinct and opposing effects on genome compartmentalization and sub compartmentalization. HP1α knockdown drives a genome-wide shift of chromatin from transcriptionally inactive B compartments to active A compartments. This is accompanied by a stepwise redistribution of A subcompartments toward the most transcriptionally active state (A3), and the upregulation of oncogenic genes involved in EMT and proliferation. In contrast, TGFβ treatment promotes chromatin compaction, increases the proportion of B compartments, and drives a stepwise reduction in active A subcompartments.

Our results highlight the differential roles of HP1α and TGFβ in shaping the 3D genome and underscore how precise, stepwise architectural changes contribute to malignant transformation. This study provides critical insight into how chromatin architecture acts as a regulatory layer in breast cancer development.

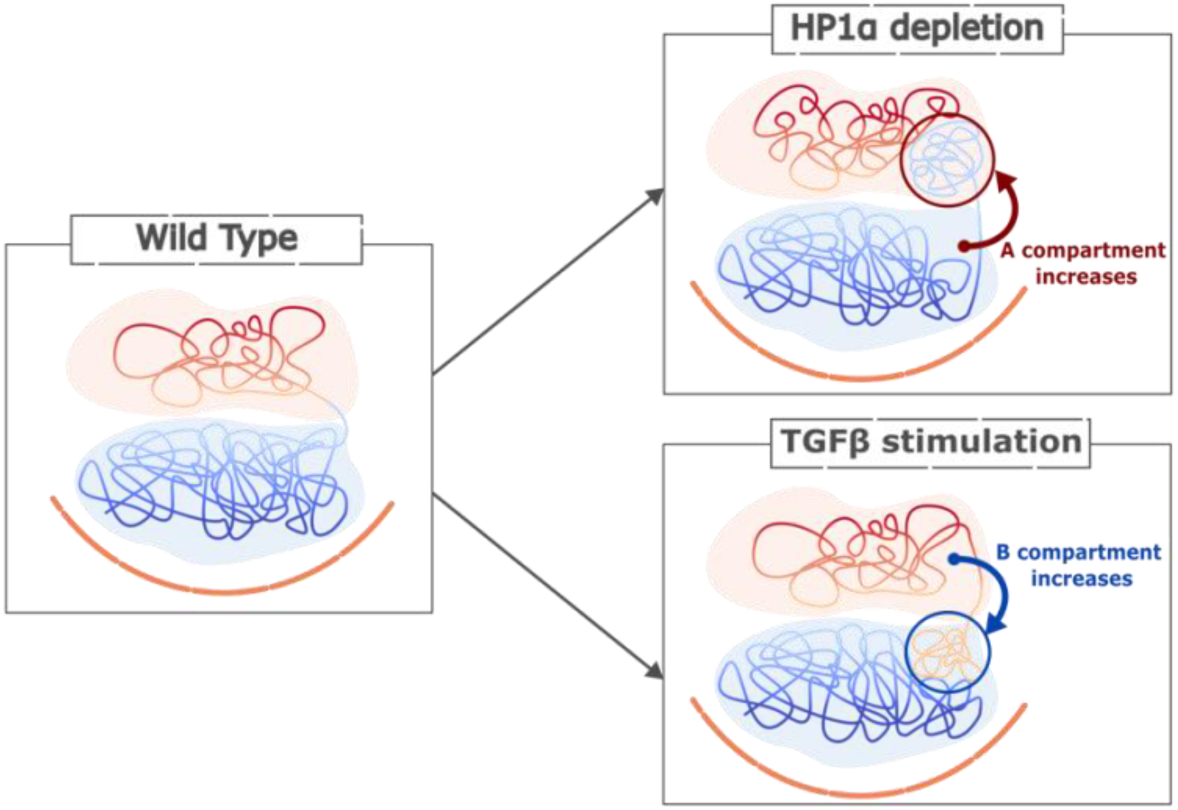
GRAPHICAL ABSTRACT.

## INTRODUCTION

The human genome is intricately organised into a hierarchical three-dimensional (3D) architecture(1), encompassing structural features such as chromatin loops (2, 3), topologically associating domains (TADs) (4, 5), larger genome compartments (6–8), and chromosome territories (9, 10). This spatial organization is not static; rather, it undergoes dynamic remodelling during key biological processes, including development and cellular differentiation (11), where it plays an important role in regulating gene expression. Disruption of this architecture is increasingly recognized as a hallmark of various pathologies, particularly in oncogenesis (12–14).

Multiple factors establish, maintain and regulate 3D genome structure. In interphase, loops and TADs are likely formed by a dynamic, transient loop extrusion process involving the Cohesin ATPase and CTCF (2, 3, 15). Compartments, on the other hand, generally organize into A and B types characterized by their association with active and repressive chromatin, respectively. A compartments are generally positioned towards the nuclear centre, while B compartments are sequestered at the nuclear periphery by chromatin interactions with lamin B receptor (LBR) and Lamin A/C (16) where they form lamina associated domains (LADs) (17). Further, intrachromosomal coalescence of compartments forms chromosomal territories where positioning of genomic regions is non-random and is correlated with transcriptional activity (10, 18). Recent studies have provided new insights into the mechanisms underlying TAD formation; however, the understanding of how higher-order structures such as genome compartments are organized, and how they are functionally linked to lower-level features, which include enhancer–promoter looping and gene regulation—remains incomplete.

Cancer development is frequently associated with profound alterations in 3D genome architecture, often coupled with widespread transcriptional dysregulation (12, 19–21). Structural features such as A/B compartments, subcompartments, and TADs have been shown to undergo extensive reorganization in breast cancer (22–24). Elucidating and comparing diverse molecular determinants, including architectural proteins e.g., HP1 (25, 26) and key players in cell proliferation and differentiation e.g., transforming growth factors (27), which govern 3D genome organization is therefore critical to understanding their roles in cancer initiation and progression.

Heterochromatin Protein 1 α (HP1α) (25, 26), a major component in establishing and regulating heterochromatin through specific binding by its chromodomain to nucleosomal histone H3 tails di– or tri-methylated on lysine 9 (H3K9me2/3) (25). This can further mediate interactions with nucleosomes and other chromatin-associated proteins (28). Upon binding to H3K9me2/3, HP1α promotes the recruitment of SUV39H, which is an enzyme that can methylate adjacent H3K9 residues, thus generating additional HP1α binding sites and thereby reinforcing heterochromatin propagation (25).

In breast cancer, the role of HP1α appears context-dependent. While some studies report that HP1α overexpression promotes cell proliferation (29), others identify it as a metastasis suppressor, with reduced expression observed in invasive breast cancer cells (30, 31). In both differentiated normal and non-metastatic breast cancer cells, HP1α contributes to the transcriptional silencing of genes involved in epithelial–mesenchymal transition (EMT), a pivotal process in metastatic progression (29, 32, 33). Supporting this, TGFβ-induced EMT in mouse mammary epithelial cells is associated with a transient reduction in HP1α protein levels during early EMT stages, coinciding with heterochromatin reorganization and acquisition of an invasive phenotype (34). Collectively, these findings implicate HP1α disruption—and the accompanying changes in 3D genome and chromatin architecture—as potentially critical drivers of the transition to metastatic breast cancer.

Transforming Growth Factor β (TGFβ) is a cytokine that signals through its receptors to regulate gene expression via SMAD family transcription factors. In humans there are three paralogous genes encoding TGF-β1, –β2, and –β3, respectively. Dysregulation of the TGFβ pathway in cancer progression involves both aberrant ligand-receptor signalling and altered transcriptional control of TGFβ genes, including autocrine loops in cells that both produce and respond to TGFβ (35). In early-stage breast cancer, TGFβ has been found to act as a tumour suppressor, however, in later stages, it promotes tumour progression (36). TGFβ signalling enhances cell mobility and activates EMT by promoting transition from epithelial to mesenchymal states (37). Furthermore, TGFβ signalling has been found to regulate promoter-super-enhancer 3D chromatin looping for TGFβ responsive genes, thus reshaping the transcriptional cellular landscape (38). It has also been shown that enhancer dynamics within TADs, with pre-existing contacts, is a main contributor to TGFβ-induced transcriptomic changes(39). Many cancer cells have been shown to express both TGFβ and its own receptors (TGFBR1 and –R2), which could result in an autocrine self-stimulating loop (40).

Despite growing evidence implicating HP1α and TGFβ in breast cancer biology, their specific roles in remodelling 3D genome architecture, and how such structural changes contribute to tumour progression, remain incompletely understood. In this study, we explore the effects of HP1α knockdown and TGFβ treatment on higher-order chromatin organization using the MCF10A human mammary epithelial cell model of breast cancer progression (41). Our findings reveal that these perturbations drive distinct and opposing alterations in chromatin sub compartmentalization: HP1α depletion shifts the genome toward a more transcriptionally permissive configuration, whereas TGFβ exposure promotes a more repressive chromatin state. These architectural changes are accompanied by dysregulation of genes involved in EMT, proliferation, and invasion, which are hallmarks of cancer progression. Together, these results highlight the differential and potentially antagonistic roles of HP1α and TGFβ in shaping the 3D genome landscape and highlights the importance of chromatin architecture as a regulatory layer in metastatic breast cancer development.

## MATERIAL AND METHODS

### Cell culture

MCF10A, and MCF10Ca1a cell lines were maintained in Dulbecco’s modified Eagle’s medium/Nutrient F12 (DMEM/F12) (Sigma-Aldrich) supplemented with 5% horse serum (MCF10A and MCF10Ca1a), 14 mM NaHCO3, 10 μg/mL insulin, 2 mM L-glutamine, 20 ng/mL human epidermal growth factor, 500 ng/mL Hydrocortisone and 100 ng/mL cholera Toxin. MCF10A cells were obtained from the American Type Culture Collection (CRL-10317). MCF10Ca1a cells were obtained from the Barbara Ann Karmanos Cancer Institute (Detroit, Michigan). HEK293T cells were maintained in Dulbecco’s modified Eagle’s medium (DMEM) (Sigma-Aldrich) supplemented with 10% v/v foetal bovine serum, 2 mM L-glutamine, and 10 U/mL penicillin and streptomycin. All cells were confirmed as mycoplasma negative.

### HP1α knockdown

To generate lentiviral particles, HEK293T cells were co-transfected using the calcium phosphate method with 20 µg of the pGFP-C-shLenti shHP1α plasmid (OriGene Technologies), 15 µg of psPAX2 (a gift from Didier Trono (Addgene plasmid # 12260; http://n2t.net/addgene:12260; RRID:Addgene_12260)), and 6 µg of pMD2.G (a gift from Didier Trono (Addgene plasmid # 12259; http://n2t.net/addgene:12259; RRID:Addgene_12259)). Viral supernatant was collected 48 hours after transfection and filtered through a 0.45 µm membrane. MCF10A cells were then transduced with the viral mixture supplemented with 4 µg/mL polybrene and 10 mM HEPES (pH 7.5). Following a 24-hour incubation, the viral medium was replaced with fresh growth media. The cells were cultured for an additional 24 hours to allow GFP expression, after which fluorescence-activated cell sorting (FACS) was performed to isolate a fully transduced cell population for downstream assays.

### TGFβ treatment

For EMT induction, MCF10A cells were treated with 5 ng/mL TGF-β1 (R&D Systems) for 8 days.

### Western Blotting

Western blot analysis was performed using standard protocols using the following antibodies: anti-HP1α (ab77256, Abcam), anti-γ-tubulin (T6557 (Clone GTU-88), Sigma-Aldrich). Blots were imaged on a Li-Cor Odyssey CLx. Protein abundance was quantified using Image Studio Lite version 5.2.5 (Li-Cor) (Supplementary Fig. S14).

### RT-qPCR

RNA was purified using the Qiagen RNeasy Plus kit and cDNA was synthesised using the Qiagen QuantiTect Reverse Transcription kit as per manufacturer’s instructions. Quantitative PCR (qPCR) was conducted on a QuantStudio 12K Flex Real-time PCR system (Life Technologies). Relative expression values were normalised to pooled housekeeping genes Glyceraldehyde 3-phosphate dehydrogenase (GAPDH), Beta-actin (β-actin), Ribosomal Protein S23 (RPS23) and Splicing Factor 3a Subunit 1 (SF3A1) (42). qPCR was performed with the following primers:

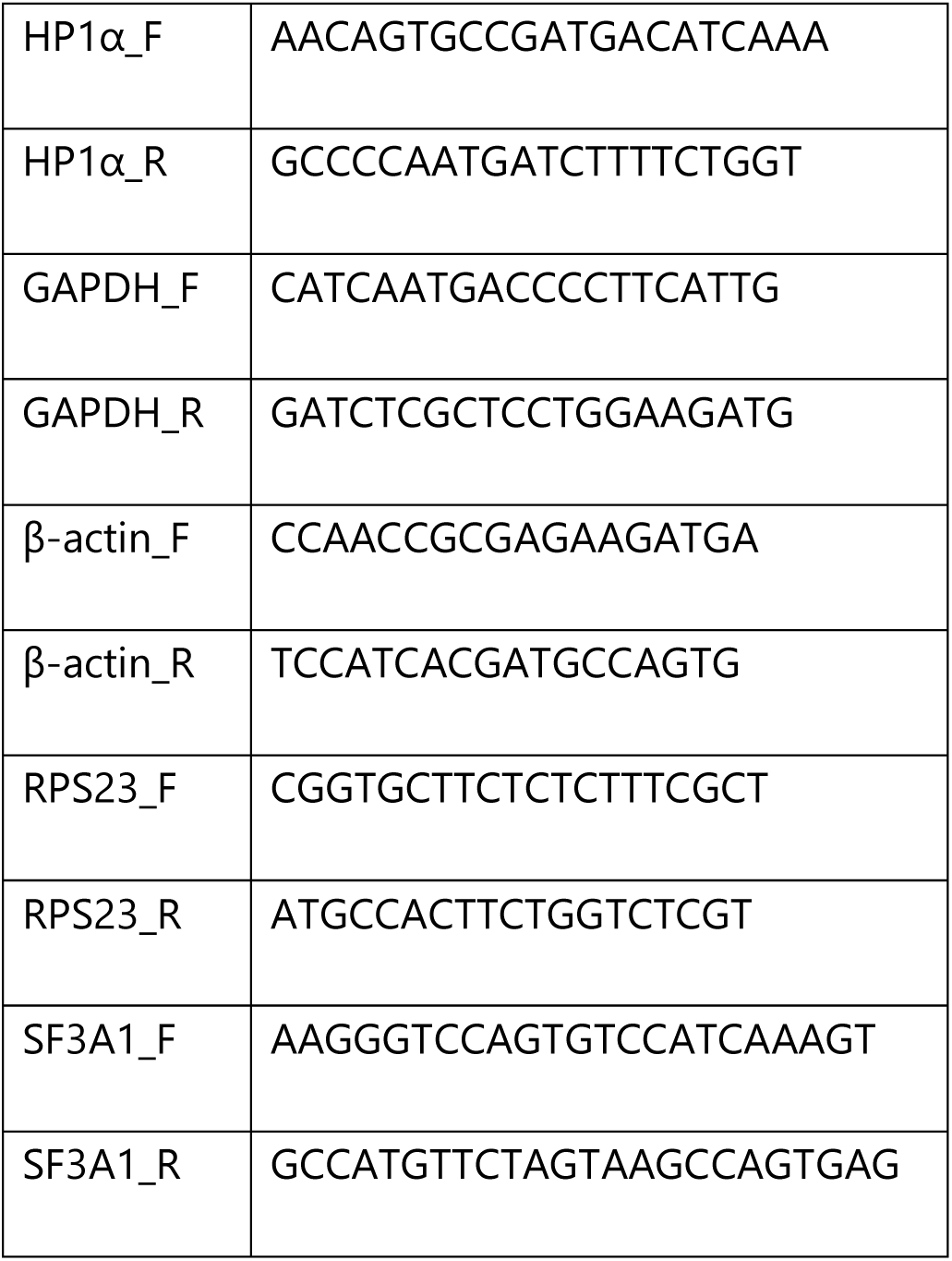

### RNA-seq

All mRNA-Seq experiments were performed in triplicate. Total RNA was isolated using the Qiagen RNeasy Plus kit following manufacturer’s instructions. Stranded mRNAseq libraries were constructed using an Illumina mRNA Prep kit according to the vendor’s protocol with poly-A enrichment. Libraries were sequenced using 76 bp paired-end reads on an Illumina NextSeq 500.

### Hi-C

Hi-C Libraries were prepared in duplicates for each cell type using the Arima-HiC+ kit (Arima Genomics, USA) according to manufacturer instructions. For each cell line, 1 million cells were used per Hi-C reaction. Cells were fixed with 1.2% formaldehyde for 12 min. All subsequent steps were carried out according to the Arima protocol. Resulting libraries were amplified with 5 PCR cycles and sequenced on an Illumina NovaSeq 6000 instrument in paired-end run with 2×101 bp.

### ChIP-seq

Approximately 1× 107 to 6 × 107 cells were cross-linked in fresh complete DMEM/F12 media supplemented with 1.2% formaldehyde at room temperature for 12 minutes. ChIP assays were carried out using anti-HP1α antibodies (ab77256, Abcam) as described previously(43). Libraries were prepared using a ChIP-Seq Sample Prep Kit (Illumina) according to the manufacturer’s protocol and sequenced on an Illumina NextSeq 500 instrument using 76 bp paired-end reads in high output mode.

### Preprocessing and analysis of Hi-C data

We used a modified version of nf-core/hic v2.1.0(44) for quality control, sequence mapping, and filtering with hg38.p14, applying the 2-enzyme Arima Kit ––restriction_site= “^GATC,G^ANTC” and –-ligation_site= “GATCGATC,GANTGATC,GANTANTC,GATCANTC”). The following flags were specified: –-skip_maps, –-skip_dist_decay, –-skip_tads, –-skip_compartments, –-skip_balancing, –-skip_mcool, –-split_fastq=false. Hi-C contact matrices in multi-resolution cooler format were generated using hictk v2.0.1 (45), and we converted the 1 kbp .cool file to .mcool using the zoomify option in hictk. Matrices were balanced using the SCALE and ICE methods for intra-chromosomal and genome-wide interactions. Downstream analyses used cis-only, balanced matrices. Biological replicates for each condition were merged using hictk merge to generate higher-depth contact maps, which were subsequently coarsened and balanced using hictk as outlined before. These merged matrices were used in specific downstream analyses, including the identification of consensus subcompartment states in genomic regions where individual replicates showed inconsistent assignments. Figure S1 was generated using hicrep v0.2.6 (46), with correlation values computed as the weighted average of individual chromosome correlation coefficients.

### Processing RNA-seq data

We utilized nf-core/rnaseq version 3.14.0 (44) to conduct quality control, trimming, alignment, and gene expression matrix generation. The pipeline was executed with the hg38.p14 reference genome and the hg38 genome annotation from Gencode v46(47). The nf-core/rnaseq pipeline was run with the following optional parameters: –-aligner=star_salmon, –-pseudo_aligner=salmon, –-extra_salmon_quant_args=’--seqBias –-gcBias’, and –-gencode=true.

### TAD analyses

We determined genomic positions of topologically associated domains (TADs) in all samples with HiCExplorer v3.7.5 (48) using default parameters on matrices at 50 kbp resolution. Supplementary Fig. S3 was created by extracting interaction data at 50-kbp resolution for each 10A TAD across multiple cell types. To minimize short-range interaction bias, contact values within 150 kbp of the diagonal were excluded. The remaining interactions were then summed and normalized based on TAD size, defined as the number of matrix bins (pixels) overlapping each TAD.

The generated TAD annotations were used to identify potential TAD fusions in the 10A.TGFB condition relative to the 10A (wild type) condition. For each 10A–10A.TGFB replicate pair, TAD borders were compared to detect overlaps. A TAD from the TGFB condition was classified as a potential fusion event if it overlapped by more than 90% with two or more distinct TADs from the 10A condition, indicating that separate domains in the wild-type may have merged into a single domain in the TGFB-treated cells. Candidate fused TADs observed consistently in at least three replicate comparisons were flagged as reproducible fusion events.

### Subcompartment analyses

Subcompartment annotations were identified using Calder v0.7 (49), which was run with default parameters. The analysis was performed using normalized Hi-C contact matrices at 10 kb. To facilitate interpretation, we reclassified Calder’s original subcompartment annotations into a simplified, numerically ordered scheme that reflects the continuum of chromatin accessibility. The four A-type subcompartments—A.1.1, A.1.2, A.2.1, and A.2.2—were relabelled as A3, A2, A1, and A0, respectively, in descending order of openness. Similarly, the B-type subcompartments—B.1.1, B.1.2, B.2.1, and B.2.2—were mapped to B0, B1, B2, and B3, corresponding to increasing levels of closedness. Based on these annotations, higher-order A/B compartments were then defined by aggregating all A-type subcompartments (A3 through A0) into a single A compartment, and all B-type subcompartments (B0 through B3) into a B compartment.

### Filtering of subcompartment switching

Building on these subcompartment annotations, we next examined regions with differing subcompartment assignments across replicates to identify a consensus subcompartment state. To identify and validate consensus subcompartment switches between replicates, we implemented a filtering procedure based on compartment identity, rank change, and switch directionality. The rank of each subcompartment is defined as a range from 0 (most closed, ’B3’) to 7 (most open, ’A3’). Switches are evaluated using three main criteria: (i) the absolute difference in rank change between replicates do not exceed 3; (ii) if the rank difference is greater than 1, switches are only considered valid if they occurred within the same compartment; and (iii) directionality of switch—defined as moving toward a more open or more closed state—have to remain consistent across replicates.

For each region with different subcompartment annotations:

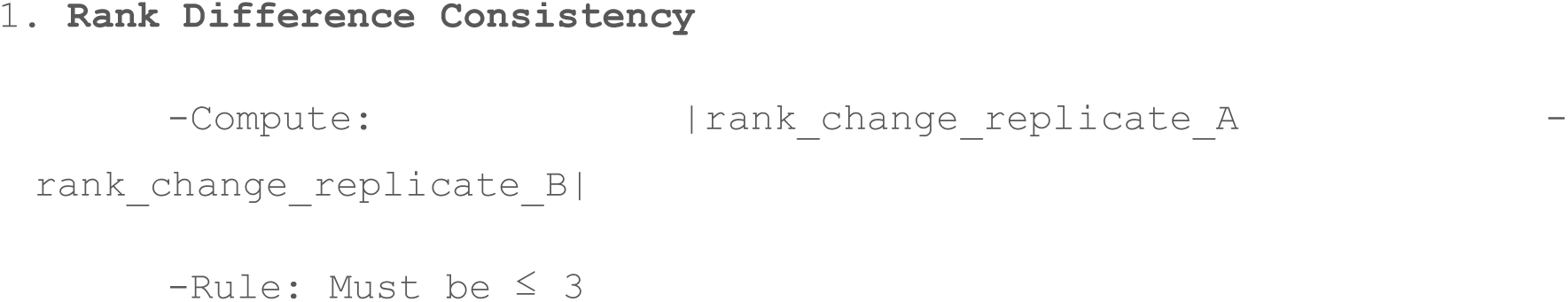
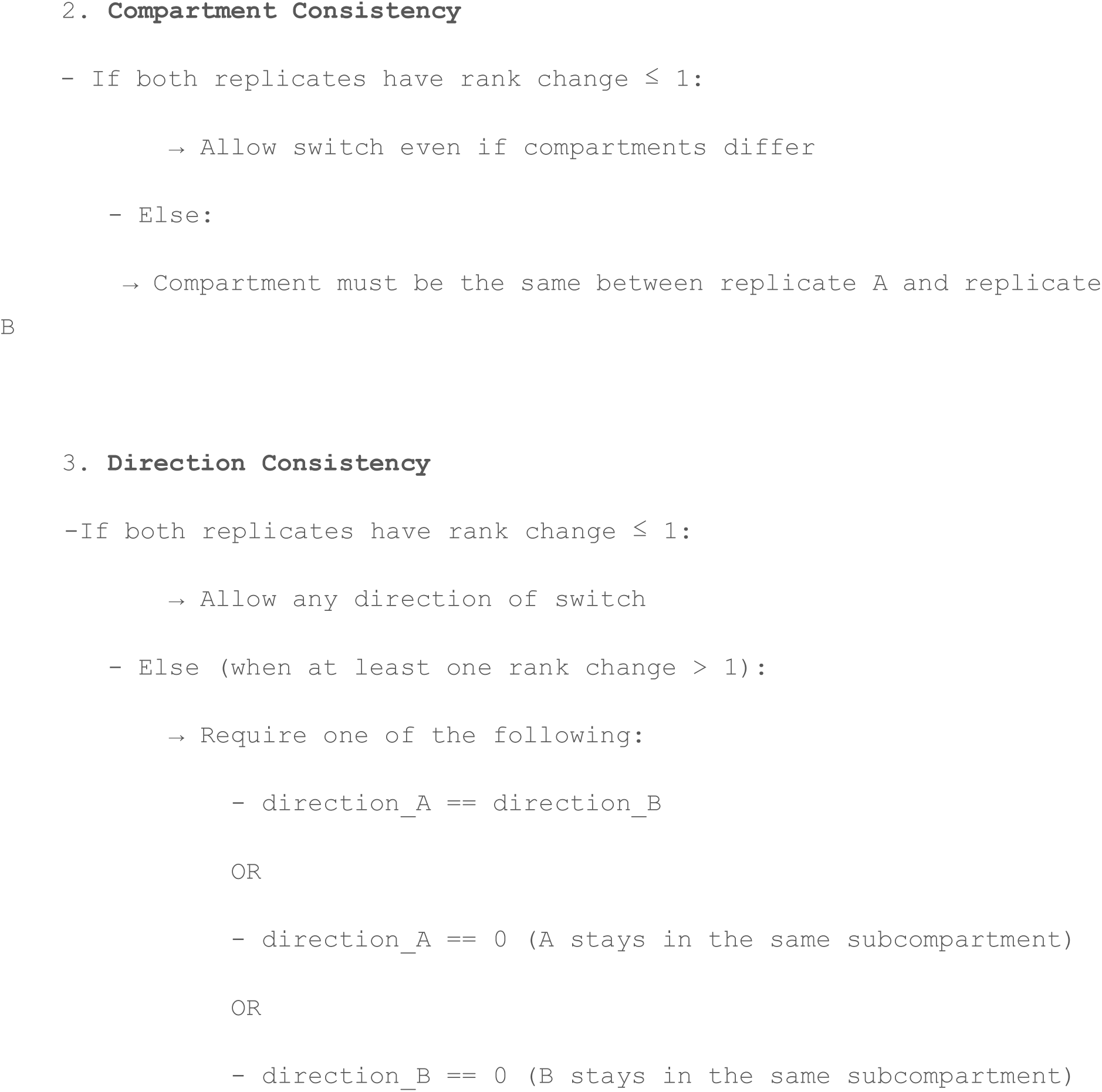

Alluvial plots were generated using the consistent switches across replicates using the R package ggalluvial(50).

### Differential Expression Analysis

Differential expression was performed using DESeq2 v1.48.0 (51) and apeglm v1.30.0 (52). DESeq2 was run with default parameters on the raw count table from nf-core/rnaseq. Log2-fold-change shrinkage was estimated with DESeq2’s lfcShrink function using apeglm and the following cutoffs: 0.0, 0.1, 0.25, 0.5, 1.0, 1.5, 2.0, 2.5, 3.0. For all the analysis the cutoff used is 0.1. Genes were considered differentially expressed if they had an absolute log2FoldChange ≥ 0.1 and a p-value ≤ 0.05 (adjusted for multiple testing using the Benjamini-Hochberg procedure).

### GO term analysis

We used genekitR v1.2.2 (53) to perform over-representation analysis and visualization of GO Biological Process terms. The IDs of switching compartment genes, subcompartments, or those included in potentially fused TADs were provided as input to genekitR. Terms were considered enriched if the adjusted p-value and q-value were both below 0.05. To simplify GO term interpretation, we utilized the GO term simplification process in genekitR. This process involves extracting species-specific GO term information from the Bioconductor organism-level annotation package for human and retrieving the ancestors of GO terms and their relationships from GO.db (54). For semantic similarity calculation, we applied the Wang algorithm from GOSemSim (55, 56).

## RESULTS

### HP1α knockdown generates open and active 3D genome compartments

To investigate the role of HP1α in 3D genome architecture, we performed duplicate Hi-C experiments in non-malignant MCF10A breast epithelial cells (“10A”) and in cells subjected to shRNA-mediated knockdown of HP1α (“10A.shHPα”) (Fig. 1A). Each sample yielded an average of ∼695 million high-quality pairwise contacts after filtering (Supplementary Table S1), enabling high-resolution analysis at 10 kbp resolution (57, 58). Quality control metrics—including the distribution of intra– (cis) and inter– (trans) chromosomal contacts—confirmed the generation of high-quality Hi-C libraries (Supplementary Table S1). Replicates were highly concordant across both conditions (SCC = 0.96–1.00; Supplementary Fig. S1), underscoring the robustness and reproducibility of the data(46). Interaction matrices were normalized using both SCALE and ICE algorithms (see Methods for details).

**Figure 1:**
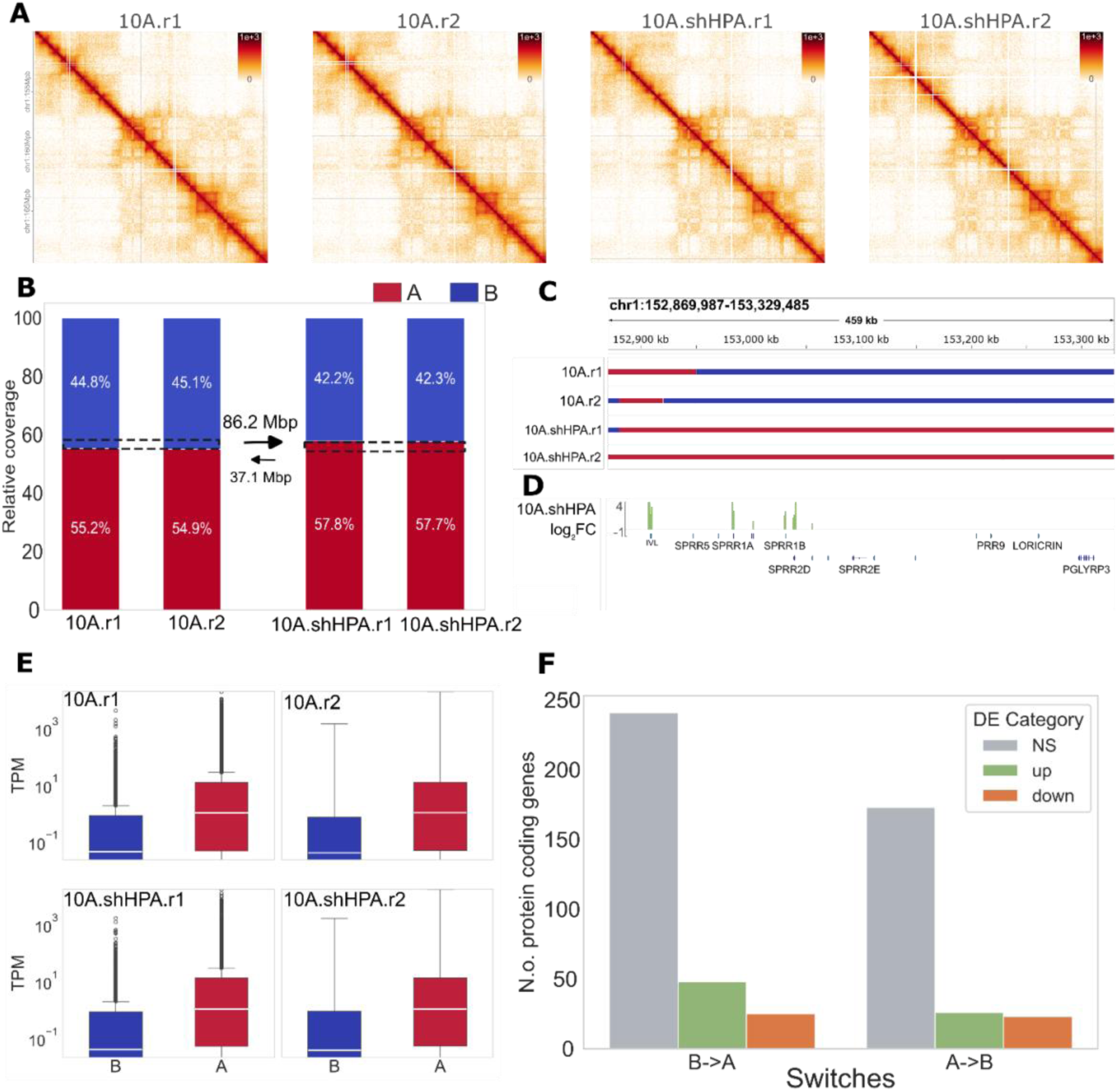
HP1α knockdown induces B-to-A compartment switching. **(A)** Hi-C contact maps at 10-kb resolution for two biological replicates of control MCF10A cells (10A) and HP1α knockdown cells (10A-shHPA). **(B)** Relative proportions of A (red) and B (blue) compartments across replicates. 10A.shHPA shows an increase in A compartment coverage, corresponding to a shift of ∼86.2 Mbp. **(C)** Genome browser view of the SPRR gene cluster (chr1:152,869,987-153,329,485), showing compartment switch from B (blue) to A (red) upon HP1α knockdown. **(D)** Genome browser view of the SPRR gene cluster (chr1:152,869,987-153,329,485) showing log2 fold change in expression (green: upregulated, orange: downregulated), calculated using a 1 kbp sliding windows. **(E)** Boxplots of gene expression levels (TPM) divided by A (red) and B (blue) compartment. **(F)** Bar plot showing the number of DEGs associated with the compartment switch. Upregulated genes are shown in green, while downregulated ones in orange. Non-significant expression changes are shown in grey.

Comparison of TADs across conditions and replicates revealed consistent features, including comparable genomic sizes (∼0.7 Mb; Supplementary Fig. S2B) and highly similar border insulation scores (R² = 0.90–1.00; Supplementary Fig. S3), indicating structural conservation of domain borders. Overall, TAD organization appeared largely unchanged between control and HP1α-depleted cells, suggesting that TAD architecture is not significantly affected after HP1α knockdown.

Given the established role of HP1α in heterochromatin maintenance and gene silencing (59, 60), we hypothesized that its depletion would induce widespread alterations in higher-order genome organization. To test this, we performed A/B compartment analysis to assess genome-wide spatial redistribution between transcriptionally active (A) and inactive (B) domains in control (MCF10A) and HP1α-knockdown (10A.shHP1α) cells. Our results revealed a notable genome-wide shift toward increased compartment A occupancy following HP1α depletion (Fig. 1B). Quantitatively, 95.12% of the genome (2404 Mbp) retained its original compartment identity (A→A or B→B), while 3.41% (86.23 Mbp) underwent a transition from B to A, and 1.47% (37.14 Mbp) from A to B (Supplementary Table S2). These data indicate that loss of HP1α causally contributes to large-scale reorganization of 3D genome architecture, promoting an overall shift toward transcriptionally permissive compartments. This indicates its direct mechanistic role in maintaining higher-order chromatin structure.

To further understand the biological relevance of these compartment shifts, we identified protein-coding genes (hereafter referred to as just genes) located within the A→B and B→A switching regions. In total, 222 genes were found in regions that switched from A to B, and 314 genes in regions that switched from B to A. The genes switching from A-to-B compartments showed no enrichment for particular biological processes. On the other hand, the genes switching from the B compartment to the A compartment were significantly enriched for biological processes including skin development and keratinization (adjusted P = 2.5e-3; Supplementary Fig. S4C). For example, a genomic region on chromosome 1 harbouring a cluster of small proline rich protein (SPRR) genes switches from B to A upon HP1α knockdown. (Fig. 1C). These genes are involved in epidermal differentiation and keratinization, and their upregulation (e.g. SPRR1A/B) are directly implicated in breast cancer (61).

To analyse gene transcriptional outcomes of HP1α knockdown, we generated triplicate RNA-seq data which were overlaid with our compartment annotations (Fig 1). As expected, genes in B compartments are less expressed than genes in A compartment (Fig. 1E). Comparing 10A and 10A.shHP1α, we identified 7,389 differentially expressed genes (DEGs), using a log2 fold-change of 0.1, among which 4,094 were upregulated and 3,295 were downregulated. To investigate whether A/B-compartment switching entails gene expression changes, we quantified whether genes that are significantly up/down-regulated are also involved in switching. For example, we noticed that the SPRR1B gene (Fig.1C) showed an upregulation (log(FC)=3.6; P=9.3×10-16) in 10A.shHPα in concordance with its switch from a B to an A compartment (Fig. 1D). Another gene, FOXA1 (a pioneer transcription factor, which plays a crucial role in development and cancer (62, 63), also switches from B to A, with an accompanying upregulation (Supplementary Fig. S5; log(FC)=0.37; P=0.002). Across the genome, genes in regions switching from B (10A) to A (10A.shHPα) compartments were significantly more frequently upregulated (P=0.0048; Binomial test), whereas genes in regions switching in the opposite direction contain a roughly equal number of up– and downregulated genes (P=0.39; Fig. 1F).

In summary, HP1α knockdown leads to the repositioning of specific genomic regions from transcriptionally repressive B compartments to active A compartments —a shift that is frequently coupled with the upregulation of genes, including those with established roles in cancer progression. These findings reinforce the function of HP1α as a chromatin-based gene silencer and suggest a causal link between its depletion, 3D genome reorganization, and specific oncogenic transcriptional activation.

### Subcompartment analysis reveals intra-compartmental switching within A compartments

Beyond the broad A/B compartment classification, the 3D genome is further partitioned into finer-scale subcompartments that reflect distinct functional and regulatory environments 50. These subcompartments are critical for interpreting how spatial genome architecture contributes to gene regulation, cellular identity, and function (64, 65). Therefore, to delve deeper into the observed HP1α-dependent compartmental alterations, we analysed our Hi-C data to determine four A sub-compartments (A0–A3) and four B sub-compartments (B0–B3) using Calder(49), where A3 represents the most transcriptionally active and B3 the most repressive subcompartment (Fig. 2A). To facilitate downstream analysis, and account for variation between replicates, we generated consensus subcompartments using a filtering and consensus-calling procedure (see Methods for details; Fig. 2B, which shows an example of inconsistent subcompartment call between the two replicates).

**Figure 2:**
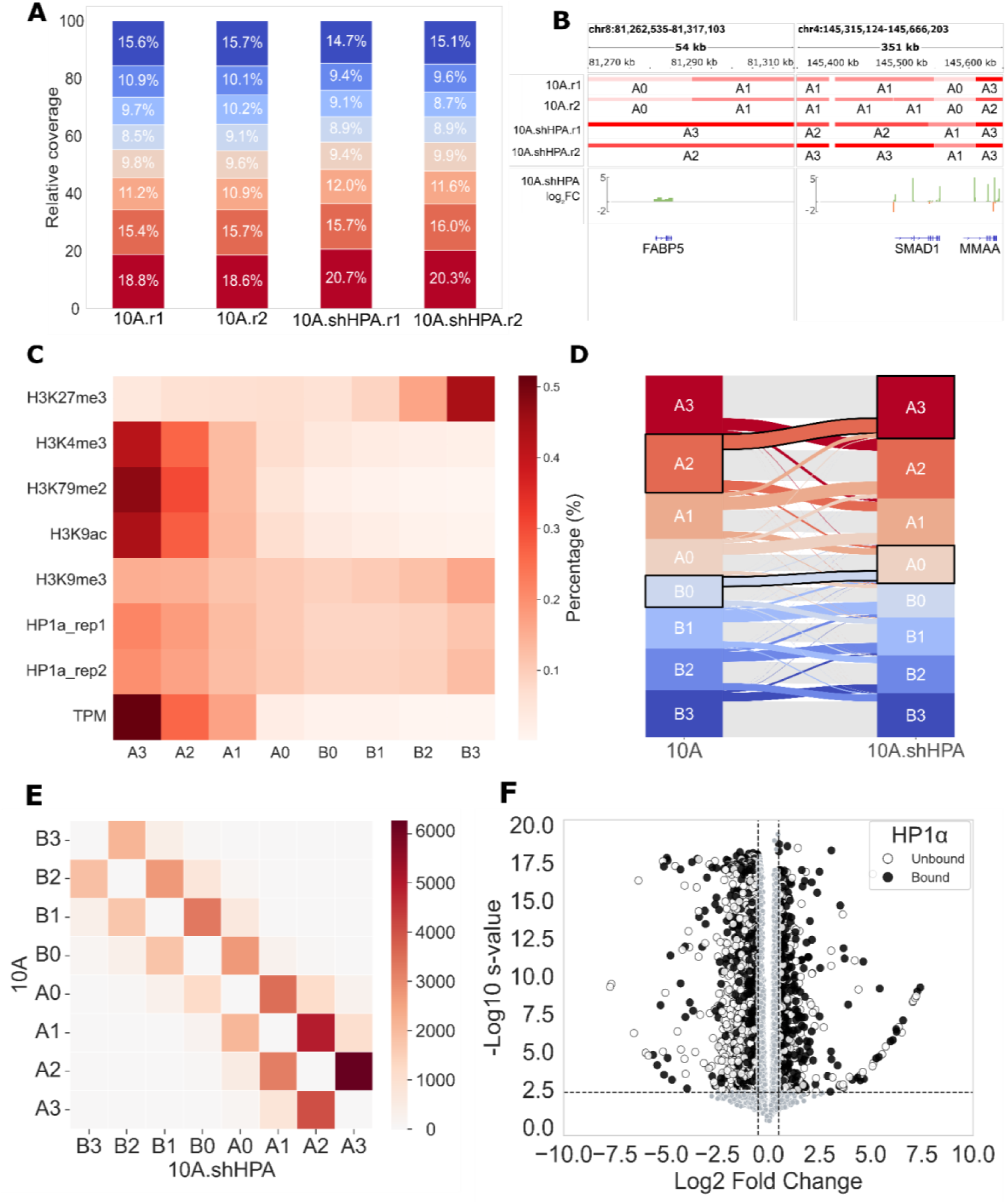
HP1a knockdown induces subcompartment reorganization favouring more open switches. **(A)** Relative genome coverage of subcompartments (10kbp resolution) across replicates. **(B)** Genome browser view of FABP5 and SMAD1 (chr8:81,262,535-81,317,103, chr4:145,315,124-145,666,203). The two genes switch to a more open chromatin state upon HP1α knockdown (10A.shHPA). These regions also show inconsistency in subcompartment calling across replicates. Tracks show gene expression log2 fold change (green: upregulated, orange: downregulated), calculated in 1 kbp sliding windows. **(C)** Heatmap showing the peak fraction of various epigenetic marks across Hi-C subcompartments in 10A.Values represent the percentage of total peaks per mark found in each subcompartment, based on raw peak counts. Signal is normalized per row to highlight relative enrichment (white: lowest, dark red: highest). **(D)** Alluvial plot of consensus subcompartment switches between 10A and 10A.shHPA. Each switch is coloured according to its subcompartment of origin. Grey lines indicate genomic regions that remain in the same subcompartment in both conditions. **(E)** Heatmap showing the number of HP1α ChIP-seq peaks associated with subcompartment switches between 10A (vertical axis) and 10A.shHPA (horizontal axis). **(F)** Volcano plot of DEGs, coloured by HP1α binding status.

Subcompartment annotations were highly consistent across replicates (Supplementary Fig. S6). Among the A sub-compartments, A0 represents the smallest fraction, comprising approximately 9.4–9.9% across all conditions, showing minimal variation between 10A and 10A.shHPA. In contrast, A3 is the largest, ranging from 18.6–20.7%, and shows an increase in 10A.shHPA, suggesting a reorganization that favours this most active subcompartment.

The B1 (∼9.7–10.2%), B2 (∼10.1–10.9%) and B3 (15.6-15.7%) all decreased slightly in 10A.shHPα, indicating some redistribution within the B sub-compartments. In contrast, B0, the smallest and least repressed B sub-compartment (8.5–9.1%), remains relatively stable (Fig. 2A). To exemplify the intercompartmental switching, the gene SMAD1 (a transcription factor, which is a central mediator in TGF-b signalling (66)) switches from A1 to A3 in 10A.shHP1a, accompanied by a statistically significant upregulation (log(FC)=0.49; P=9.48×10-19) (Fig. 2B, right panel). The gene FABP5 (a key regulator of cell metabolism and growth that is overexpressed in cancers (67)), switches from A0 to A3, also with a significant upregulation (log(FC)=0.80; P=2.18×10-80). The observed increase in A compartment occupancy following HP1α depletion aligns with HP1α’s established function in heterochromatin formation and maintenance(25).

Overlaying publicly available histone modification profile data revealed distinct chromatin features across subcompartments. A3 subcompartments are enriched in active chromatin marks (H3K4me3, H3K79me2, and H3K9ac) and gene expression (TPM) and depleted for the repressive mark (H3K27me3). The B-subcompartment, on the other hand, shows an opposite trend moving gradually from B0 towards B3 (Fig. 2C). Thus, our subcompartments represent a continuous, yet distinct chromatin state annotation from the most active (A3) towards the least active (B3) parts of the genome.

To better understand which genomic regions are affected by HP1α knockdown, we performed ChIP-seq for HP1α in 10A cell samples. Overlaying the resulting binding peaks on the subcompartments revealed that HP1α shows chromatin binding in all subcompartments, yet with a slight preference for A3 (19.3% of the peaks) and B3 (12.95%) subcompartments (percentages based on the peak count, Fig. 2C). The slight preference of HP1α for A subcompartments was initially surprising but suggested that it has a fine-tuning transcriptional role at active genes (see below).

Investigating an alluvial plot of switching events between subcompartments (Fig. 2D), reveals that the increase in A3 observed under knockdown conditions (Fig. 2A) appears to result primarily from switching between consecutive subcompartments, rather than direct shifts from B subcompartments to A3. Notably, B-to-A switches are predominantly driven by B0 regions moving into A0 (Fig. 2D (highlighted box; ∼59 Mbp, 2.3%). This initial shift is accompanied by extensive reorganization within both the A and B subcompartments. Among the A subcompartments, a substantial fraction of A2 regions further switches into A3 (Fig. 2D; highlighted box; 113 Mbp; 4.5%), contributing to its overall enrichment. While switches within both A and B subcompartments are bidirectional, they are generally biased toward more open subcompartments, consistent with a global shift toward more open A-subcompartments following HP1α knockdown (Fig. 2D). Regions switching to a more open state are enriched for GO terms related to various immune functions and sodium ion transmembrane transport (Fig. S7A).

We next examined whether the observed subcompartment switches were enriched in regions binding HP1α, based on our ChIP-seq data. To assess this, we analysed HP1α peak enrichments in relation to all subcompartment switching events by statistically comparing HP1α peak enrichments in each switch direction compared to a switch in opposite direction (Fig. 2E, Fig. S8). This revealed an asymmetric switching pattern, with certain switches showing a significant enrichment with respect to the opposite switch. Notably, genomic regions switching from B0 to A0 (2,723 vs. 1,192 peaks; P ≈ 7 × 10-136) showed significantly higher HP1α enrichment compared to the reverse switch. Similar trends were observed for switches from A2 to A3 (6,261 vs. 4,057 peaks; P ≈ 2.16 × 10-105) and B1 to A0 (678 vs. 334 peaks; P ≈ 6.5 × 10-28). Overall, binomial testing revealed that HP1α preferentially associates with genomic regions shifting toward more open subcompartments upon the knockdown (see Supplementary Table S3). We also investigated genes within switching regions together with the expression changes. This analysis revealed a consistent pattern: genes located in regions switching towards a more open subcompartment were more likely to be upregulated. Notably, genes within B0 to A0 were among the most enriched for upregulation (29 vs. 10, P ≈ 0.002), aligning with the observed HP1α binding enrichment. Similarly, A2 to A3 switches represented the most significant category associated with upregulated genes (330 vs. 193, P ≈ 1.1e-09, Supplementary Table S4).

Finally, we examined how HP1α binding at the TSS of genes relates to their differential gene expression upon HP1α knockdown. Volcano plots of all genes indicate that binding of HP1α is more prevalent at the TSS of significantly upregulated (2,353 genes of 4,094 upregulated genes) compared to downregulated genes (1,433 of 3,295 downregulated genes). Using Fisher’s exact test, this difference is statistically significant (P=5.8 x 10-33; Supplementary Table S5), indicating that genes bound by HP1α are more likely to be upregulated upon HP1α knockdown (Fig. 2F). Taken together, these results show: (1) HP1α exhibits unexpected binding in active chromatin; (2) subcompartment switching is stepwise and biased toward openness; (3) HP1α binding correlates with regions prone to opening and (4) Genes with HP1α at the TSS are preferentially upregulated upon knockdown.

### TGFβ stimulation oppositely restructures 3D genome towards B compartments

In addition to HP1α, dysregulation of TGFβ signalling is a well-established driver of invasive breast cancer progression(68–71). TGFβ promotes EMT, a fundamental process underlying tumour invasiveness and metastasis(70, 72, 73). Importantly, both TGFβ and HP1α have been shown to regulate EMT through distinct yet possibly interacting chromatin-based mechanisms(31, 33). To explore this further, we investigated how TGFβ stimulation and HP1α depletion differentially or similarly affect higher-order chromatin architecture. To this end, we generated replicate Hi-C datasets from MCF10A cells treated with TGFβ (hereafter referred to as “10A.TGFB”) and compared the resulting chromatin organization with that observed upon HP1α knockdown (Fig. 3A).

**Figure 3:**
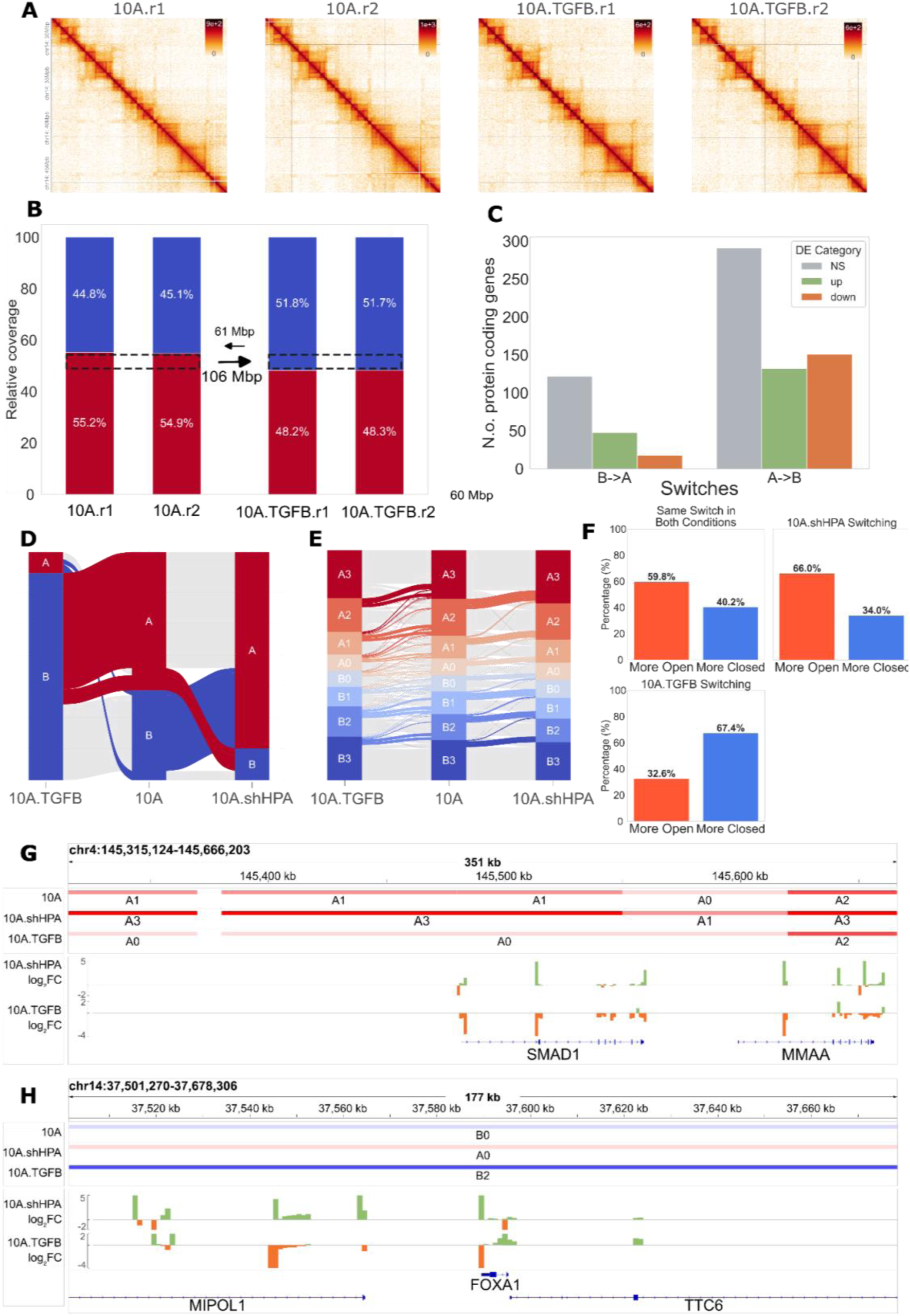
TGFβ stimulation promotes closed compartment, with subcompartment rewiring opposite to HP1α’s direction. **(A)** Hi-C contact maps at 10-kb resolution for two biological replicates of control MCF10A cells (10A) and TGFβ treated cells (10A-TGFB). **(B)** Relative proportions of A (red) and B (blue) compartments across replicates. **(C)** Bar plot showing the number of DEGs associated with compartment switches in 10A.TGFB. Upregulated genes are shown in green, while downregulated ones in orange. Non-significant expression changes are shown in grey. **(D)** Alluvial plot showing a zoomed view of A/B compartment switches, restricted to regions that undergo switching in either TGFβ-treated (10A.TGFB), HP1α knockdown (10A.shHP1α), or both. **(E)** Alluvial plot showing switches to a more open state from 10A to 10A.shHPA and to a more closed state from 10A to 10A.TGFB. **(F)** Bar plots quantifying the directionality of subcompartment switches, categorized as shared or condition-specific based on whether the same switch occurs across conditions or uniquely in one. **(G)** Genome browser view of the SMAD1 (chr8:81,262,535-81,317,103), showing consensus subcompartments for 10A, 10A.shHPA and 10A.TGFB. Bottom tracks show gene expression log2 fold change (green: upregulated, orange: downregulated), calculated in 1 kbp sliding windows. **(H)** Genome browser view of the FOXA1 (chr14:37,501,270-37,678,306), showing consensus subcompartments for 10A, 10A.shHPA and 10A.TGFB. Bottom tracks show gene expression log2 fold change (green: upregulated, orange: downregulated), calculated in 1 kbp sliding windows.

Quality control metrics confirmed high data quality and reproducibility, with strong correlation between replicates (SCC = 0.96–1.00; Supplementary Fig. S1), consistent TAD sizes (∼0.7 Mbp), and highly concordant insulation scores (R² = 0.90–1.00; Supplementary Fig. S3). Notably, despite the overall preservation of TAD architecture, we observed a subtle reduction in the number of TADs in TGFβ-treated samples compared to controls, with an average loss of ∼75 TADs (Supplementary Fig. S2B). The observed reduction in TAD number can be attributed to TAD fusion events, though technical differences cannot be ruled out (Supplementary Fig. S9A). Across replicates, we identified ∼120 consistent fused TADs spanning approximately 150 Mbp of the genome (see Method for more details). Within these, there are ∼900 genes which show enrichment for processes such as negative regulation of growth and detoxification of inorganic compounds (Supplementary Fig. S9B).

In contrast to HP1α knockdown, which promotes chromatin opening, TGFβ treatment induces a genome-wide shift toward a more closed chromatin state, such that the proportion of B compartments increased from 44.8–45.1% in control (10A) cells to 51.7–51.8% in 10A.TGFB (Fig. 3B). The total switching is ∼167 Mbp, with 106.36 Mbp (4.27%) switching from the A compartment into the B compartment. 60.96 Mbp (2.45%) switches from the B (10A) to the A compartment (10A.TGFB) (Supplementary Table S6). Genes located in regions that switched from A (10A) to B (10A.TGFB) compartments were significantly more likely to be downregulated (P = 0.0001, Binomial test). In contrast, while a higher number of upregulated genes was observed in regions switching from B to A, this trend was not statistically significant (P = 0.88, Binomial test) (Fig 3C).

To assess the relationship between compartment changes induced by TGFβ treatment and those resulting from HP1α knockdown, we visualized compartment switches across 10A, 10A.TGFB, and 10A.shHP1α cells using alluvial diagrams (Fig. 3D). This analysis revealed limited overlap, with 49.38 Mbp of shared compartment switches: 19 Mbp from A to B and 30 Mbp from B to A. We further examined the GO terms associated with genes found in switching regions. Consistent with compartment data, there was a limited overlap in enriched GO terms between 10A.TGFB and 10A.shHPA. Notably the only common term common to both treatments in switching compartment genes was “defense response to Gram-positive bacterium”, associated in both cases with regions switching from B to A (Fig. S10).

We further explored chromatin changes by generating consensus subcompartment annotations and tracking their redistribution across conditions (Fig. 3E, S11, S12). In 10A.TGFB cells, we observed a marked reduction in A2 and A3 subcompartments (from 18.8–18.6% to 16.6–16.6% for A3, and from 15.4–15.7% to 13.2–13% to A2), coupled with a shift of A0/A1 regions into the B compartment (99 Mbp, 93% of the total switches). Interestingly, 10A.shHP1α cells displayed a similar reorganization within the A compartment, but in the opposite direction, with switching favouring more transcriptionally active subcompartments (Fig. 3E).

Of all subcompartment switches, 16% were unique to TGFβ-treated cells, 10.8% were specific to HP1α knockdown, and 18.9% were shared (Fig. S13). Among these shared switches, similar fractions appeared towards more open (59.8%) and closed (40.2%) compartments (Fig. 3F; top left panel). A statistically significant majority of TGFβ-specific switches moved toward a closed chromatin state rather than a more open state (67.4% and 32.6%, respectively; P<1e-50 [Binomial test]; Fig.3F bottom left panel), whereas HP1α-specific switches were significantly enriched for compartmental opening rather than closing (66% and 34%, respectively; P<1e-50 [Binomial test]; Fig. 3F top right panel). For example, the genomic region harbouring genes SMAD1 switches from an A1 subcompartment in 10A to a A0 subcompartment in 10A.TGFB cells, and switches to A3 in 10A.shHPA (Fig. 3G). The gene FOXA1 is found in a B0 subcompartment in 10A, but switches to an A0 compartment in 10A.shHPA but becomes B2 in 10A.TGFB (Fig. 3H).

Collectively, these results reveal: (1) TGFβ promotes a genome-wide shift toward a more repressive chromatin state; (2) Compartment switching analysis revealed limited overlap between TGFβ-induced and HP1α-depletion-induced chromatin changes indicating that TGFβ and HP1α act largely through independent chromatin remodelling pathways; (3) TGFβ treatment produces a distinct chromatin reorganization compared to HP1a depletion highlighted by their contrasting effects on active subcompartment switching and (4) Key genes implicated in cancer, such as SMAD1 and FOXA1, display differential subcompartment localization depending on the treatment.

### Both HP1α knockdown and TGFβ stimulation show dysregulation of genes implicated **in breast cancer development**

Given that both HP1α and TGFβ have been implicated in breast cancer progression, we next compared their transcriptional effects to gene expression patterns observed during different stages of tumorigenesis. Specifically, we analysed differential gene expression (DEG) profiles from malignant (MCF10ACa1a, hereafter “10A.C1”) stages of the MCF10A breast cancer progression model(12). HP1α expression was significantly reduced in the malignant C1 stage (Fig. 4A). Surprisingly, TGFβ expression also showed downregulation in C1 (Fig. 4B), despite its known role in promoting EMT and invasion.

**Figure 4:**
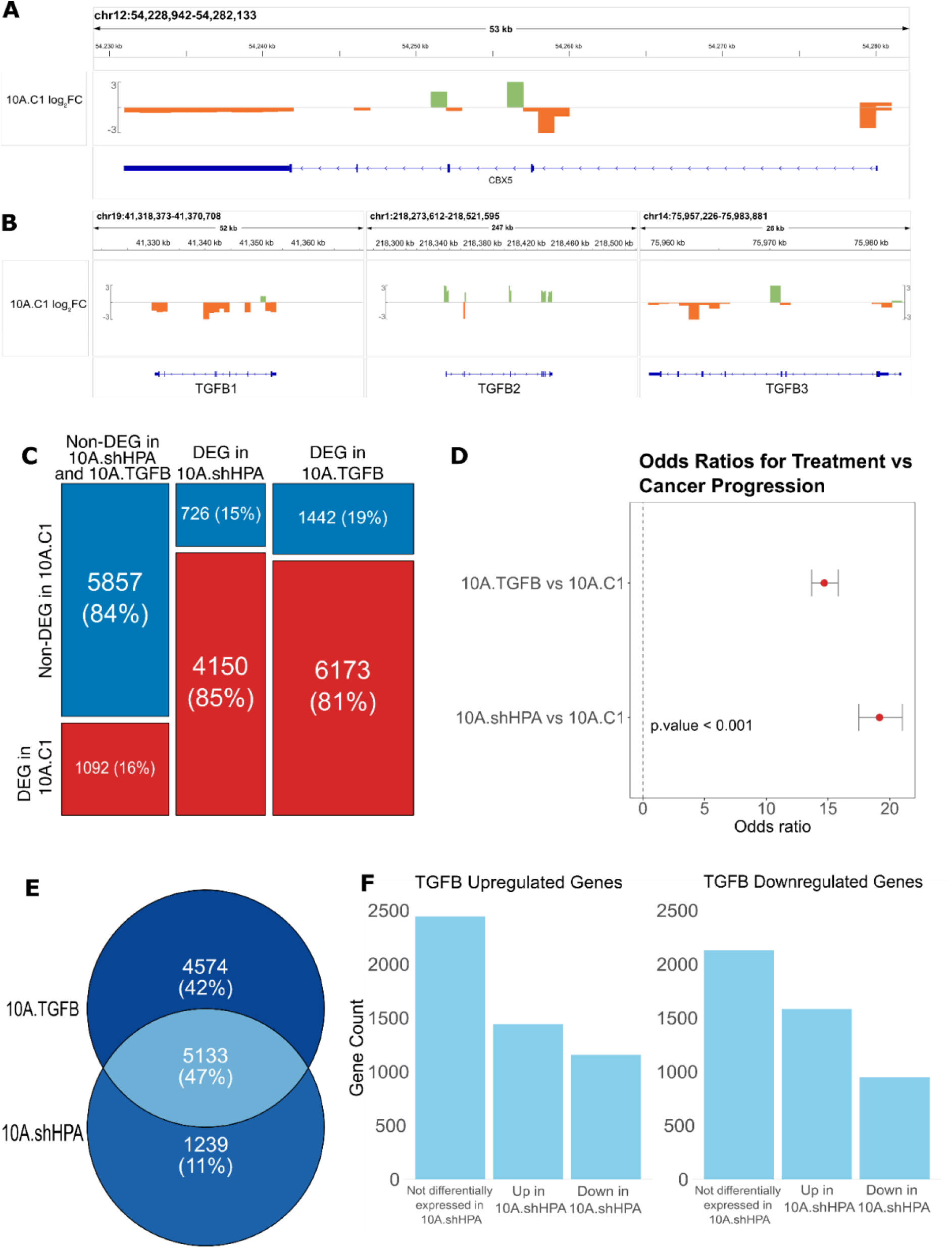
HP1α knockdown and TGFβ treatment resemble gene expression profiles of cancer progression. **(A)** Genome browser view of CBX5 (HP1α) (chr12:54,228,942-54,282,133), showing gene expression log2 fold change (green: upregulated, orange: downregulated), calculated in 1 kbp sliding windows in 10A.C1 compared with 10A **(B)** Genome browser view of TGFB1 (chr19:41,318,373-41,370,708), TGFB2 (chr1:218,273,612-218,521,595) and TGFB3 (chr14:75,957,226-75,983,881) showing gene expression log2 fold change (green: upregulated, orange: downregulated), calculated in 1 kbp sliding windows in 10A.C1. **(C)** Overlap of DEGs between treatment conditions (10A.TGFB and 10A.shHP1a) and malignant cells (10A.C1). In the plot, blue indicates genes that are not differentially expressed in 10A.C1, red represents DEGs specific to 10A.C1. **(D)** Odds ratio analysis comparing treatment-induced DEGs with gene expression changes observed in malignant cells (10A.C1). Red represents DEGs specific to 10A.C1. **(E)** Venn diagram showing the overlap of differentially expressed genes (DEGs) between 10A.TGFB and 10A.shHP1a treatments, restricted to the subset of genes that are also differentially expressed in the 10A.C1. **(F)** 10A.TGFB differentially expressed genes overlap with upregulated and downregulated genes in 10A.shHPA.

To determine the extent to which the transcriptional responses induced by HP1α knockdown and TGFβ stimulation resemble those occurring during tumour progression, we compared the DEGs from each treatment with those identified in the C1 progression stages. This revealed a substantial overlap: 5,729 genes (54%) were shared between 10A.shHP1α and 10A.C1 (Fig. 4C). Chi-square tests confirmed that these overlaps were highly significant (p < 0.001), indicating that the similarities are unlikely to arise by chance. This suggests that both HP1α depletion and TGFβ stimulation induce gene expression changes aligned with the oncogenic reprogramming (Fig. 4D). To further examine whether these changes are coordinated, we analysed the overlap in DEGs between the two treatments within the subset of genes also differentially expressed in the malignant cells. Approximately 47% of DEGs in 10A.TGFB were also altered in 10A.shHP1α (Fig. 4E), with the majority showing upregulation in response to HP1α knockdown (Fig. 4F).

In conclusion, these results show that: (1) Both HP1α knockdown and TGFβ treatment trigger transcriptional changes that closely mirror early-stage breast cancer gene expression; (2) These perturbations show biologically meaningful overlap with gene expression programs associated with oncogenic transformation and (3) While HP1α knockdown and TGFβ stimulation differ in their effects on 3D genome structure, their shared transcriptional outcomes highlight a convergence on early cancer-related gene networks.

## DISCUSSION

Recent advances in cancer biology have underscored the importance of 3D genome organization and nuclear architecture in tumour formation and progression(12, 14). Alterations in chromatin structure, particularly within compartment and subcompartment domains, are increasingly recognized as key contributors to malignant transformation(12). However, the mechanisms by which chromatin remodelling and signalling pathways converge to alter genome topology and gene regulation during cancer progression remain poorly defined.

Although HP1α is classically associated with heterochromatin and transcriptional repression, its role in 3D genome organization and its contribution to breast cancer development have not been fully elucidated. In this study, we systematically dissect how HP1α depletion and TGFβ signalling influence genome architecture and transcriptional states in a breast epithelial cell model. Importantly, our work provides mechanistic insight into how these factors impact compartmental organization, sub compartmental redistribution, and oncogenic gene expression programs.

Our findings highlight the critical role of HP1α in maintaining a repressive 3D genome architecture and regulating transcription through compartmental organization. Upon HP1α knockdown, we observed a genome-wide redistribution of chromatin from transcriptionally inactive B compartments toward active A compartments, consistent with a more open and permissive chromatin state (Fig. 1B; Supplementary Table S2). In contrast, TGFβ treatment induced the opposite effect, driving an increase in B compartment occupancy and a more repressive chromatin environment (Fig. 3B; Supplementary Table S4), aligning with previous studies reporting TGFβ-mediated chromatin repression(39).

Functional analysis of regions switching from B to A in 10A.shHP1α cells revealed a significant enrichment for genes involved in epidermal development and keratinization (adjusted P = 2.5e-3; Supplementary Fig. S5C), biological processes frequently co-opted during tumour progression and EMT. Notably, this included SPRR1A and SPRR1B, genes known to promote aggressive tumour phenotypes by enhancing proliferation, resistance to therapy, and cell motility(61). A particularly noteworthy example is FOXA1, a pioneer transcription factor essential for breast tissue differentiation. In HP1α-depleted cells, FOXA1 undergoes a compartment switch into a more transcriptionally active state and is concurrently upregulated. FOXA1 is capable of binding compacted chromatin and facilitating its remodelling, thereby enabling transcription factors such as estrogen receptor α (ERα) to access target loci(62). In breast cancer, aberrant overexpression of FOXA1 has been linked to increased invasiveness and endocrine resistance by reshaping the ER-dependent transcriptional landscape(62, 67, 74). These results illustrate how HP1α loss not only remodels genome architecture but also activates key regulatory genes associated with malignant transformation.

Conversely, TGFβ stimulation induces the opposite pattern-promoting chromatin compaction and increasing the proportion of B compartments. This global shift toward transcriptionally repressive chromatin is associated with the downregulation of genes residing in A→B switching regions, consistent with TGFβ’s role in driving epithelial–mesenchymal transition (EMT) and metastatic behaviour through gene repression(39, 70, 72, 73, 75). Beyond broad compartmental changes, subcompartment analysis reveals that both HP1α knockdown and TGFβ treatment elicit extensive reorganization within the A subcompartments, but also in functionally opposing directions. HP1α depletion promotes redistribution into A3 —the most transcriptionally active subcompartment— while TGFβ drives a reduction in A2 and A3 subcompartments, reflecting a transition toward a more repressed chromatin state. These shifts are largely driven by stepwise transitions between adjacent subcompartments, rather than abrupt compartmental switching, highlighting a ‘fine-tuned’ restructuring of genome structure. Genes such as FABP5 and SMAD1 exemplify this pattern, both transitioning toward A3 and becoming upregulated following HP1α knockdown. Notably, SMAD1 has emerging roles in EMT and metastasis through interactions with MEK/ERK signalling(74, 76, 77).

ChIP-seq profiling of HP1α revealed that regions most likely to switch toward more active chromatin states were pre-marked by HP1α binding (Fig. 2E). For example, transitions such as B0→A0 and A2→A3 showed significant HP1α enrichment prior to knockdown. Genes with HP1α occupancy at their transcription start sites were also significantly more likely to be upregulated upon knockdown, supporting a model in which HP1α directly dampens active promoters (Fig. 2F).

Collectively, this study identifies HP1α as a causal regulator of compartmental and sub compartmental integrity in breast epithelial cells. Its depletion drives stepwise chromatin opening and transcriptional activation of EMT– and cancer-related genes. In contrast, TGFβ signalling, while similarly promoting EMT and invasion, induces a global shift toward chromatin compaction and transcriptional repression. These findings highlight two distinct, yet complementary, modes of chromatin-based regulation converging on shared oncogenic programs. Understanding how structural genome dynamics integrate signalling and epigenetic cues offers new insights into the molecular basis of tumour initiation and provides potential avenues for therapeutic intervention.

## ACKNOWLEDGEMENTS

Not applicable

## AUTHOR CONTRIBUTIONS

JP, RA and DT conceived and designed the study; FP processed Hi-C and RNA-seq data and performed all downstream analyses, including interpretation of results; YD performed cell culture work and RNA-seq; MN did HiC experiments; BA established the HP1 knockdown cell lines; TS performed the HP1 Chip-Seq; OH drafted the introduction and contributed to the results section; JP, RA, FP drafted the manuscript, with inputs from DT and OH. JP, RA, DT supervised the work. All authors read and approved the final manuscript.

## SUPPLEMENTARY DATA STATEMENT

Supplementary Data are available at NAR online.

## DATA AVAILABILITY

All raw and processed sequencing data generated in this study have been submitted to the NCBI Gene Expression Omnibus (GEO; https://www.ncbi.nlm.nih.gov/geo/) under accession numbers GSE316327. The code used for the data analysis is available at 10.5281/zenodo.18465946 (78).

