## supplemental data for "HP1α depletion and TGFβ activation exert antagonistic effects on 3D genome organization"

**Table S1**: Hi-C interactions by sample.

| | 10A.rep1 | 10A.rep2 | 10A.shHPA.rep1 | 10A.shHPA.rep2 | 10A.TGFB.rep1 | 10A.TGFB.rep2 |
|---|---|---|---|---|---|---|
| **Valid Pairs** | 634.4 | 723.1 | 627.1 | 693.7 | 760.8 | 736.6 |
| **Read Pairs** | 735.5 | 864.1 | 739.9 | 797.6 | 917.7 | 882 |
| **cis interaction** | 419.9 | 478.6 | 404 | 447.1 | 462.5 | 446.2 |
| **cis interaction (shortRange)** | 111.4 | 126.7 | 123.7 | 136.4 | 897.2 | 846.7 |
| **trans interaction** | 103.2 | 117.8 | 99.4 | 110.2 | 208.5 | 205.7 |
| **cis trans ratio** | 4.07 | 4.06 | 4.06 | 4.06 | 2.22 | 2.17 |
| **cis long-short ratio** | 0.26 | 0.26 | 0.31 | 0.31 | 1.94 | 1.90 |

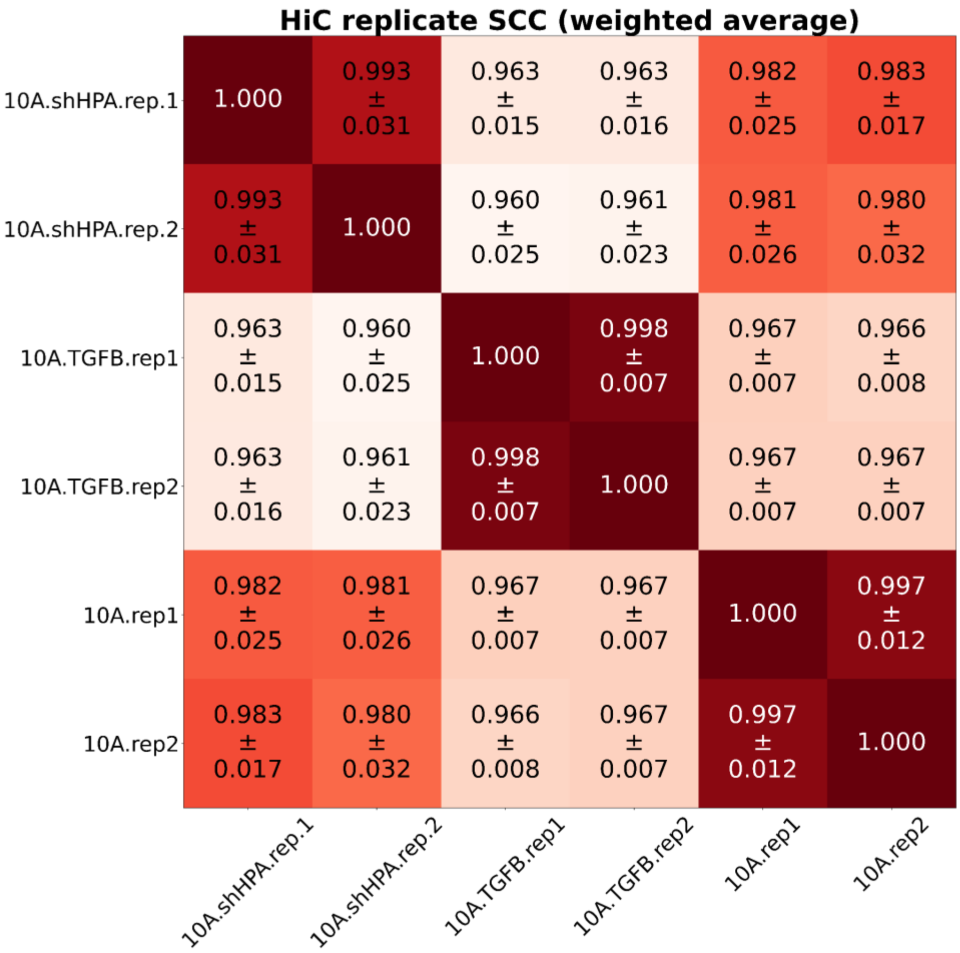

**Figure S1**: Heatmap displaying the weighted average of stratified correlation coefficients (SCC) calculated for each sample pair utilizing HiCRep. This method accounts for the distance-dependent decay characteristic of HiC data to prevent overestimated correlations. The computation of the weighted average SCC incorporates chromosome sizes as weights. The values represent the weighted average along with the standard deviation for each sample pair.

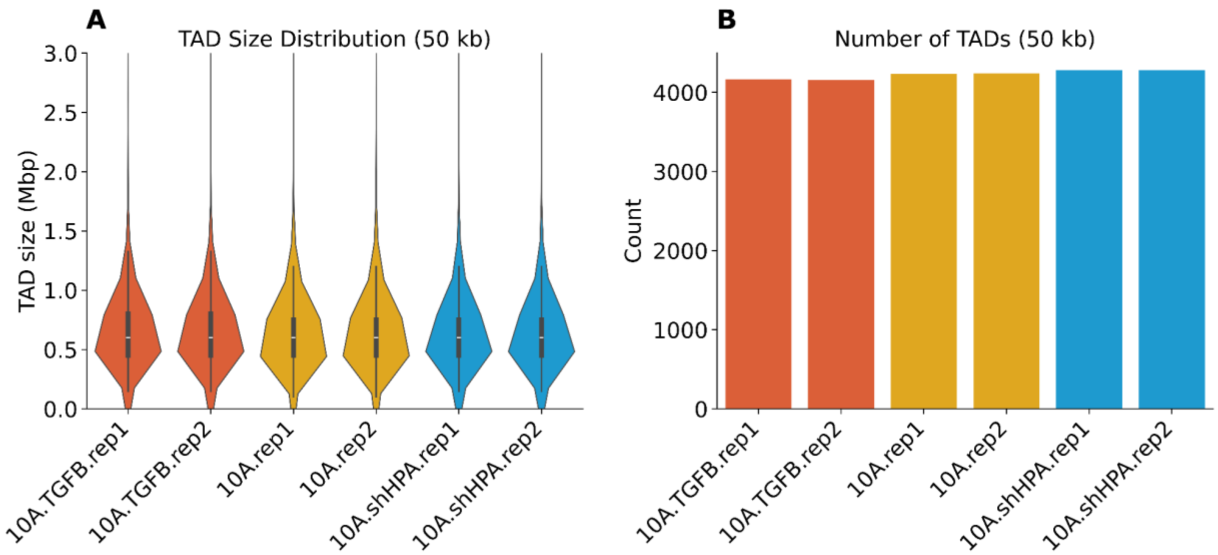

**Figure S2**: **A** Total number of TADs identified in each of the six generated Hi-C samples.
**B** Genomic size distribution of all TADs across the six samples. The TAD sizes are generally comparable among the samples, though a slight reduction in the number of TADs is noticeable in the TGFβ-treated samples.

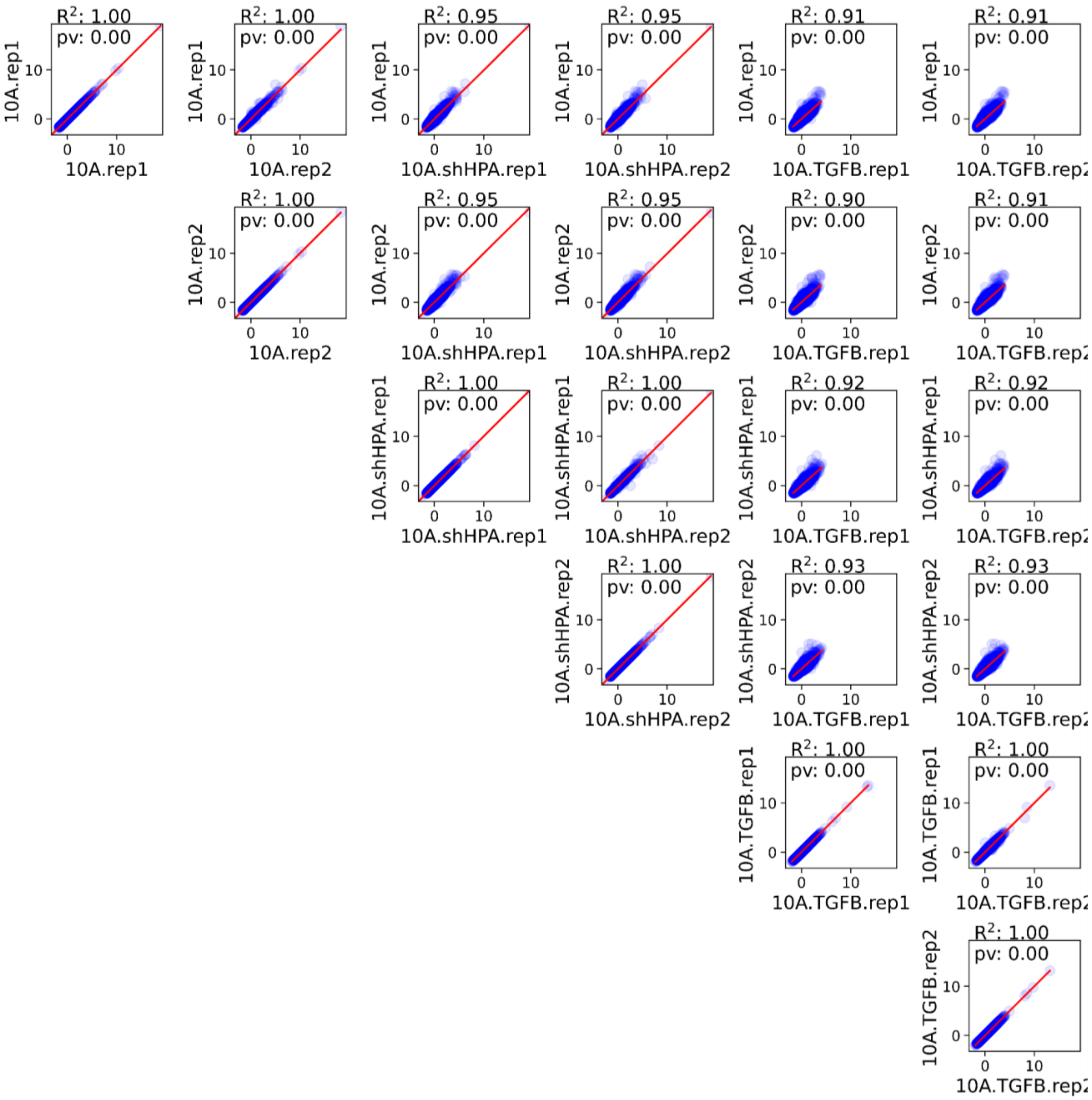

**Figure S3**: Scatterplot showing a comparison of TAD insulation scores at a 50 kbp resolution for every pairwise combination of samples. The red line represents the observed trend.

**Table S2**: Coverage of compartment switches in 10A.shHPA

| 10A | 10A.shHPA | Coverage (Mbp) | Relative Coverage (%) |
|---|---|---|---|
| A | A | 1363.3 | 53.94 |
| A | B | 37.14 | 1.47 |
| B | A | 86.23 | 3.41 |
| B | B | 1040.7 | 41.18 |

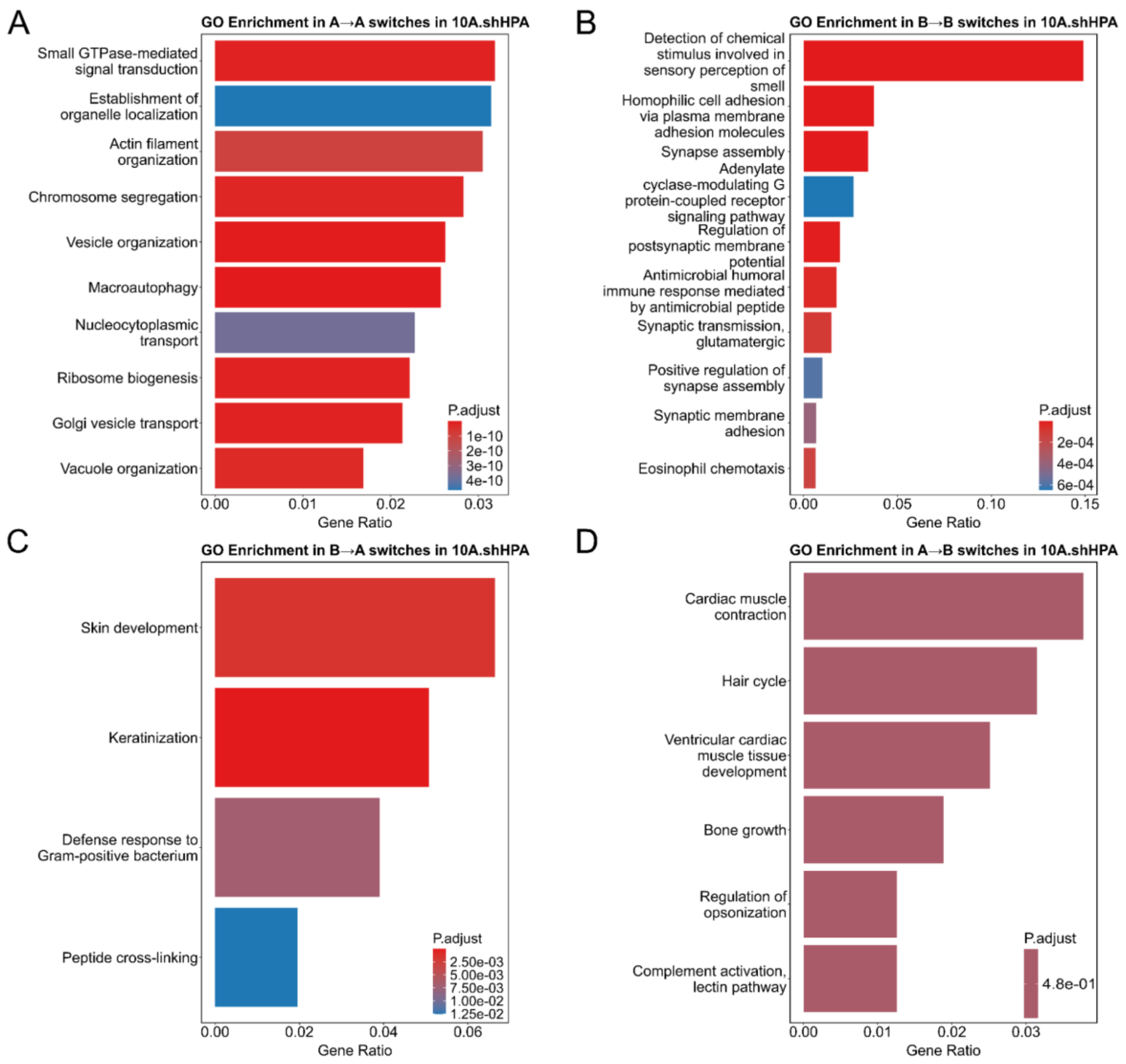

**Figure S4**: **A** GO enrichment analysis in regions switching from A (10A) to A (10A-shHPA) compartment,
**B** GO enrichment analysis in regions switching from B (10A) to B (10A-shHPA) compartment,
**C** GO enrichment analysis in regions switching from B (10A) to A (10A-shHPA) compartment,
**D** GO enrichment analysis in regions switching from A (10A) to B (10A-shHPA) compartment.

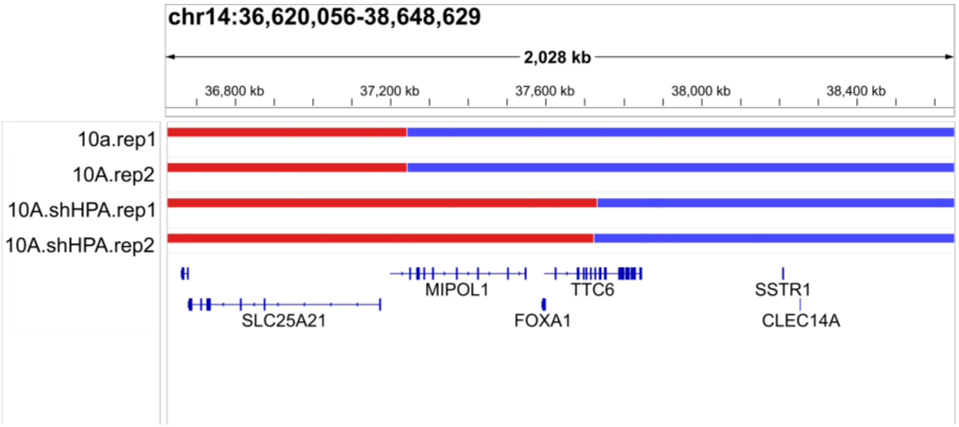

**Figure S5**: Genome browser view of chromosome 14, 36,620,056-38,648,629, showing a compartment switch from B (blue) to A (red) upon HP1α knockdown.

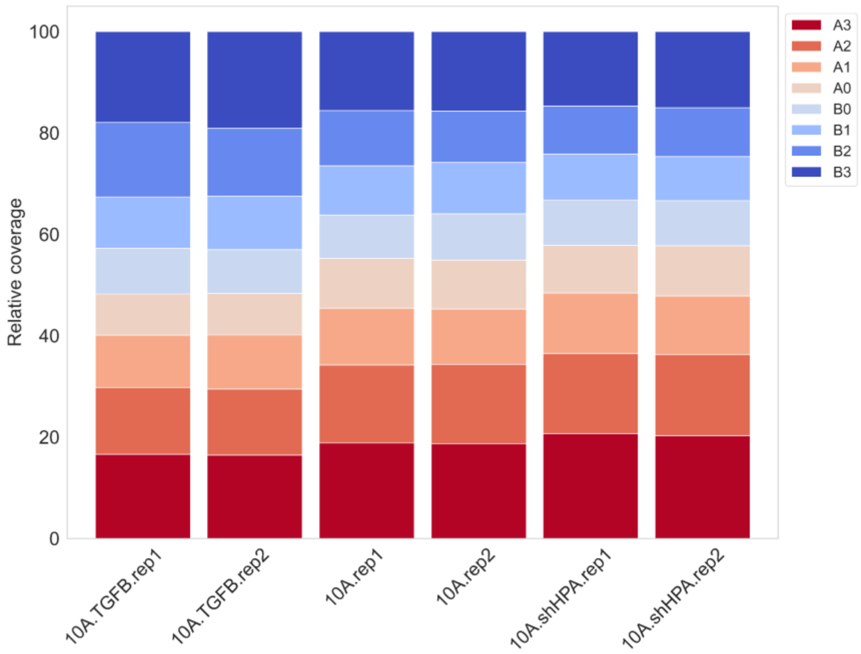

**Figure S6** Relative genome coverage of subcompartments (at 10kbp resolution) in each replicate for each condition.

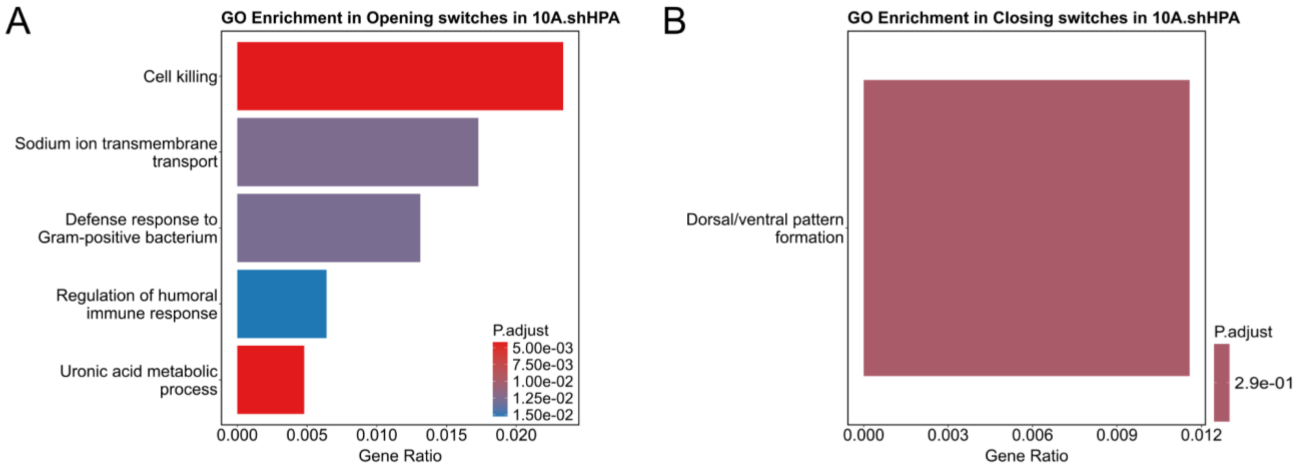

**Figure S7**: **A** GO enrichment analysis in regions switching from a more closed compartment in 10A to a more open compartment in 10A-shHPA

**B** GO enrichment analysis in regions switching from a more open compartment in 10A to a more closed compartment in 10A-shHPA

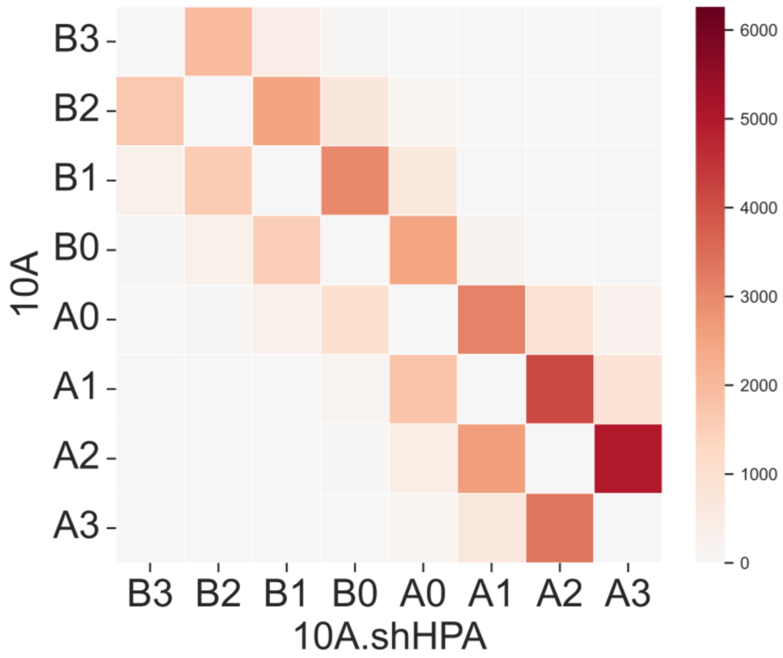

**Figure S8**: Heatmap showing the number of HP1α ChIP-seq peaks associated with subcompartment switches between 10A and 10A.shHPA.

**Table S3**: Binomial test p-values for HP1α peaks for all pairwise subcompartment switches.

| from | to | count_from_to | count_to_from | ratio | p_value |
|---|---|---|---|---|---|
| B0 | A0 | 2723 | 1192 | 0.7 | 7.0e-136 |
| A2 | A3 | 6261 | 4057 | 0.61 | 2.2e-105 |
| B1 | B0 | 3286 | 1768 | 0.65 | 7.1e-103 |
| A0 | A1 | 3516 | 2088 | 0.63 | 3.6e-82 |
| A1 | A2 | 4892 | 3181 | 0.61 | 1.2e-81 |
| A0 | A2 | 1132 | 509 | 0.69 | 8.2e-55 |
| B2 | B1 | 2705 | 1740 | 0.61 | 4.8e-48 |
| B2 | B0 | 753 | 382 | 0.66 | 7.6e-29 |
| B1 | A0 | 678 | 334 | 0.67 | 6.5e-28 |
| A0 | A3 | 260 | 142 | 0.65 | 2.1e-09 |
| A1 | A3 | 1057 | 824 | 0.56 | 4.3e-08 |
| B2 | A1 | 29 | 3 | 0.91 | 1.3e-06 |
| B2 | A0 | 122 | 61 | 0.67 | 3.8e-06 |
| A3 | B2 | 15 | 0 | 1.0 | 3.5e-05 |
| B3 | B2 | 2109 | 1863 | 0.53 | 5.0e-05 |
| A3 | B0 | 13 | 1 | 0.93 | 0.0009 |

| | | | | | |
|---|---|---|---|---|---|
| A2 | B0 | 84 | 49 | 0.63 | 0.001 |
| B3 | B1 | 433 | 370 | 0.54 | 0.01 |
| B3 | A0 | 14 | 4 | 0.78 | 0.01 |
| A3 | B3 | 6 | 0 | 1.0 | 0.02 |
| A2 | B3 | 6 | 0 | 1.0 | 0.02 |
| B0 | B3 | 99 | 73 | 0.58 | 0.03 |
| B0 | A1 | 252 | 214 | 0.54 | 0.04 |
| B1 | A2 | 14 | 7 | 0.67 | 0.09 |
| B2 | A2 | 10 | 7 | 0.59 | 0.31 |
| B1 | A1 | 22 | 20 | 0.52 | 0.44 |
| B1 | A3 | 1 | 0 | 1.0 | 0.5 |
| B3 | A1 | 1 | 0 | 1.0 | 0.5 |
| A1 | B1 | 20 | 22 | 0.48 | 0.68 |
| A2 | B2 | 7 | 10 | 0.41 | 0.83 |
| A2 | B1 | 7 | 14 | 0.33 | 0.96 |
| A1 | B0 | 214 | 252 | 0.46 | 0.96 |
| B3 | B0 | 73 | 99 | 0.42 | 0.98 |
| B1 | B3 | 370 | 433 | 0.46 | 0.98 |

| | | | | | |
|---|---|---|---|---|---|
| A0 | B3 | 4 | 14 | 0.22 | 0.996 |
| B0 | A2 | 49 | 84 | 0.37 | 0.999 |
| B0 | A3 | 1 | 13 | 0.07 | 1.0 |
| B2 | B3 | 1863 | 2109 | 0.47 | 1.0 |
| A0 | B2 | 61 | 122 | 0.33 | 1.0 |
| A1 | B2 | 3 | 29 | 0.09 | 1.0 |
| A3 | A1 | 824 | 1057 | 0.44 | 1.0 |
| A3 | A0 | 142 | 260 | 0.35 | 1.0 |
| A1 | B3 | 0 | 1 | 0.0 | 1.0 |
| B3 | A2 | 0 | 6 | 0.0 | 1.0 |
| A3 | B1 | 0 | 1 | 0.0 | 1.0 |
| B3 | A3 | 0 | 6 | 0.0 | 1.0 |
| B2 | A3 | 0 | 15 | 0.0 | 1.0 |
| A2 | A0 | 509 | 1132 | 0.31 | 1.0 |
| B1 | B2 | 1740 | 2705 | 0.39 | 1.0 |
| B0 | B2 | 382 | 753 | 0.34 | 1.0 |
| B0 | B1 | 1768 | 3286 | 0.35 | 1.0 |
| A0 | B1 | 334 | 678 | 0.33 | 1.0 |

| | | | | | |
|---|---|---|---|---|---|
| A1 | A0 | 2088 | 3516 | 0.37 | 1.0 |
| A0 | B0 | 1192 | 2723 | 0.3 | 1.0 |
| A2 | A1 | 3181 | 4892 | 0.39 | 1.0 |
| A3 | A2 | 4057 | 6261 | 0.39 | 1.0 |

**Table S4:** Binomial test p-values for upregulated genes for all pairwise subcompartment switches.

| from | to | count_from_to | count_to_from | ratio | p_value |
|---|---|---|---|---|---|
| A2 | A3 | 330 | 193 | 0.63 | 1.1e-09 |
| A1 | A2 | 178 | 93 | 0.66 | 1.3e-07 |
| A0 | A1 | 85 | 44 | 0.66 | 0.0002 |
| B0 | A0 | 29 | 10 | 0.74 | 0.002 |
| A0 | A2 | 25 | 11 | 0.69 | 0.01 |
| A1 | A3 | 32 | 24 | 0.57 | 0.17 |
| A0 | A3 | 9 | 5 | 0.64 | 0.21 |
| B3 | B0 | 3 | 1 | 0.75 | 0.31 |
| B1 | B3 | 4 | 2 | 0.67 | 0.34 |
| B1 | A0 | 4 | 2 | 0.67 | 0.34 |
| B2 | A0 | 4 | 2 | 0.67 | 0.34 |

| | | | | | |
|---|---|---|---|---|---|
| B3 | B2 | 5 | 3 | 0.62 | 0.36 |
| B1 | B0 | 19 | 18 | 0.51 | 0.5 |
| A2 | B0 | 2 | 1 | 0.67 | 0.5 |
| B3 | A1 | 1 | 0 | 1.0 | 0.5 |
| B2 | A1 | 1 | 0 | 1.0 | 0.5 |
| B1 | B2 | 7 | 7 | 0.5 | 0.6 |
| B2 | B1 | 7 | 7 | 0.5 | 0.6 |
| B0 | B1 | 18 | 19 | 0.49 | 0.6 |
| B0 | A1 | 3 | 3 | 0.5 | 0.7 |
| A1 | B0 | 3 | 3 | 0.5 | 0.7 |
| B1 | A1 | 1 | 1 | 0.5 | 0.75 |
| B0 | B2 | 1 | 1 | 0.5 | 0.75 |
| B2 | B0 | 1 | 1 | 0.5 | 0.75 |
| A1 | B1 | 1 | 1 | 0.5 | 0.75 |
| B2 | B3 | 3 | 5 | 0.38 | 0.85 |
| B0 | A2 | 1 | 2 | 0.33 | 0.88 |
| A3 | A1 | 24 | 32 | 0.43 | 0.89 |
| A0 | B1 | 2 | 4 | 0.33 | 0.89 |

| | | | | | |
|---|---|---|---|---|---|
| A0 | B2 | 2 | 4 | 0.33 | 0.89 |
| B3 | B1 | 2 | 4 | 0.33 | 0.89 |
| A3 | A0 | 5 | 9 | 0.36 | 0.91 |
| B0 | B3 | 1 | 3 | 0.25 | 0.94 |
| A2 | A0 | 11 | 25 | 0.31 | 1.0 |
| A0 | B0 | 10 | 29 | 0.26 | 1.0 |
| A1 | A0 | 44 | 85 | 0.34 | 1.0 |
| A2 | A1 | 93 | 178 | 0.34 | 1.0 |
| A3 | A2 | 193 | 330 | 0.37 | 1.0 |
| A1 | B2 | 0 | 1 | 0.0 | 1.0 |
| A1 | B3 | 0 | 1 | 0.0 | 1.0 |
| B3 | A0 | 0 | 0 | | |
| B3 | A2 | 0 | 0 | | |
| B3 | A3 | 0 | 0 | | |
| B2 | A2 | 0 | 0 | | |
| B2 | A3 | 0 | 0 | | |
| B1 | A2 | 0 | 0 | | |
| B1 | A3 | 0 | 0 | | |

| | | | |
|---|---|---|---|
| B0 | A3 | 0 | 0 |
| A0 | B3 | 0 | 0 |
| A2 | B3 | 0 | 0 |
| A2 | B2 | 0 | 0 |
| A2 | B1 | 0 | 0 |
| A3 | B3 | 0 | 0 |
| A3 | B2 | 0 | 0 |
| A3 | B1 | 0 | 0 |
| A3 | B0 | 0 | 0 |

**Table S5:** Contingency table showing the association between HP1α binding at gene TSS and differential gene expression upon HP1α knockdown

| | **HP1α Bound** | **HP1α Not Bound** |
|---|---|---|
| **Upregulated** | 2353 | 1741 |
| **Downregulated** | 1433 | 1862 |

**Table S6**: Coverage of compartment switches in 10A.TGFB

| 10A | 10A.shHPA | Coverage (Mbp) | Relative Coverage (%) |
|---|---|---|---|
| A | A | 1237.73 | 49.74 |
| A | B | 106.36 | 4.27 |
| B | A | 60.96 | 2.45 |
| B | B | 1083.37 | 43.54 |

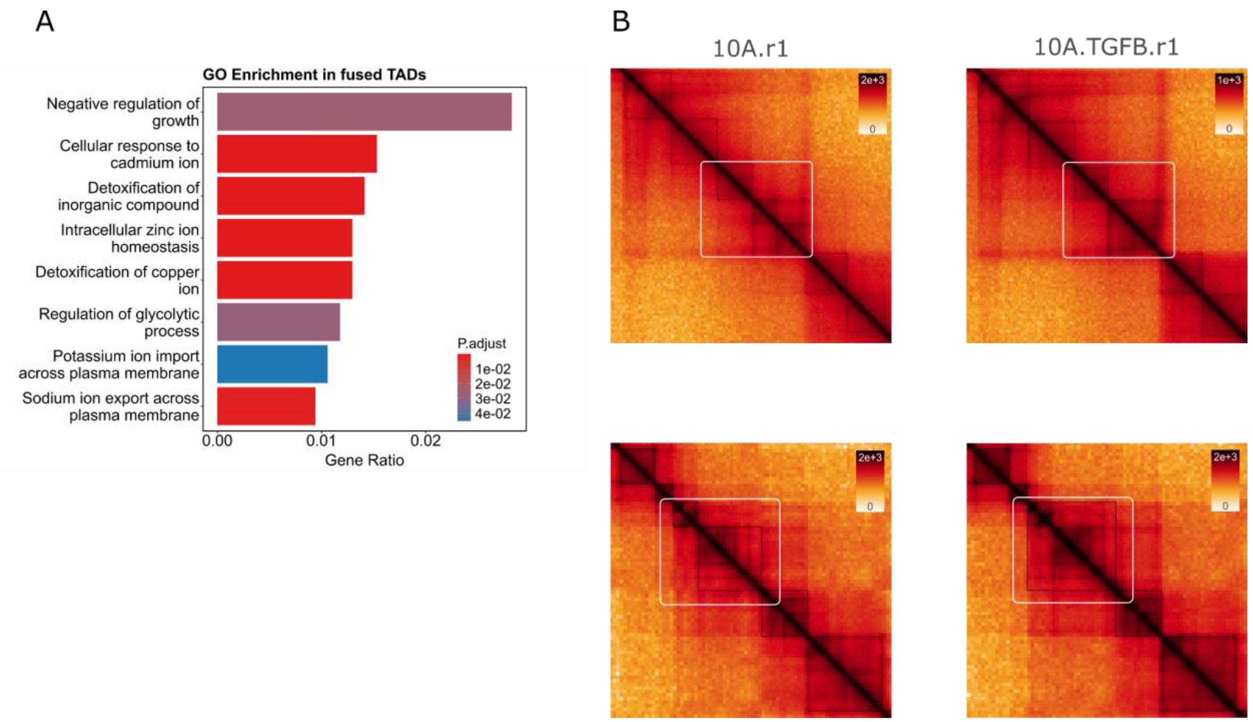

**Figure S9**: **A** GO enrichment analysis for genes located in putatively fused TADs,
**B** Contact maps showing a representative biological TAD fusion (top) and a technical fusion resulting from boundary-calling artifacts (bottom). White squares highlight the fused regions in both instances.

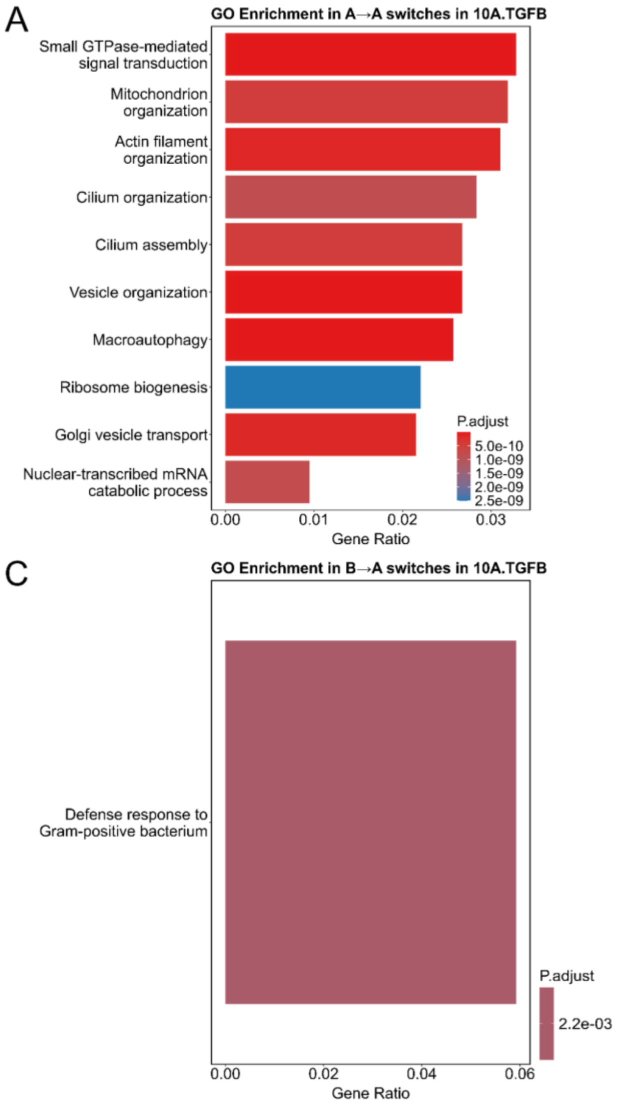

**Figure S10**: **A** GO enrichment analysis in regions switching from A (10A) to A (10A-TGFB) compartment,
**B** GO enrichment analysis in regions switching from B (10A) to B (10A-TGFB) compartment,
**C** GO enrichment analysis in regions switching from B (10A) to A (10A-TGFB) compartment,
**D** GO enrichment analysis in regions switching from A (10A) to B (10A-TGFB) compartment.

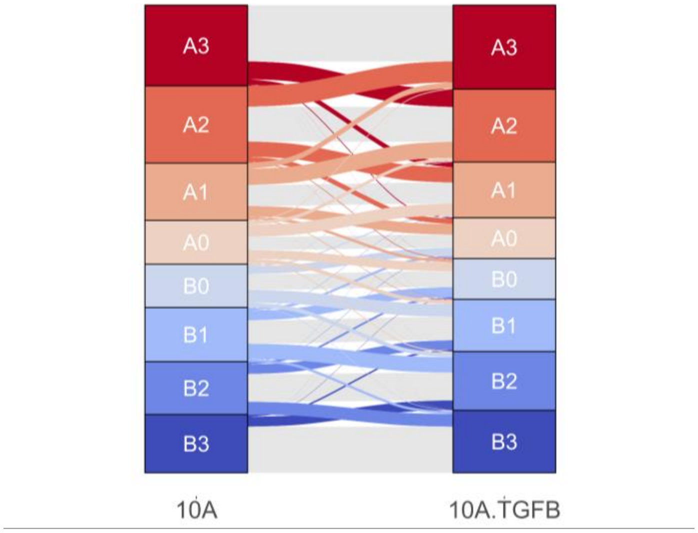

**Figure S11**: Alluvial plot showing switches from 10A to 10A.TGFB.

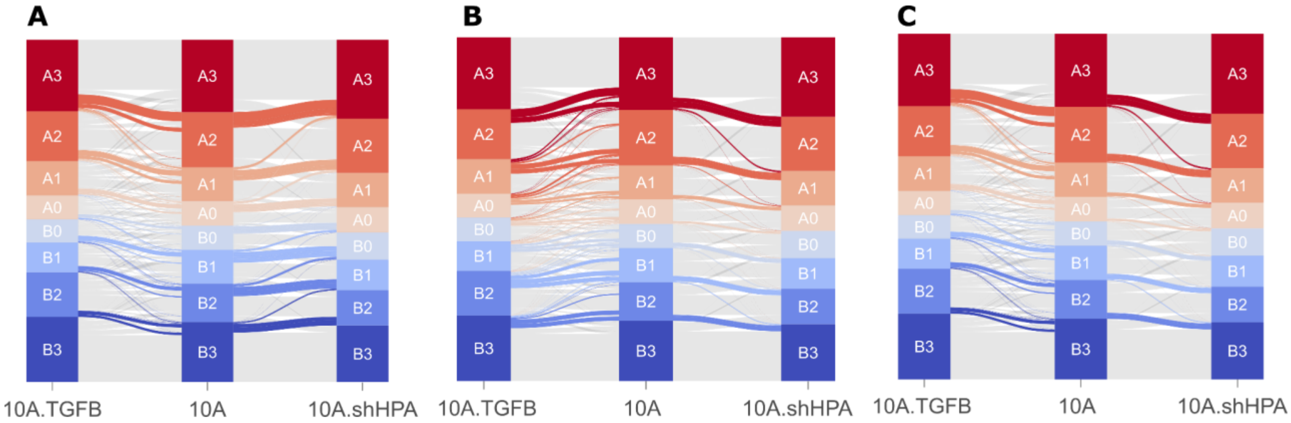

**Figure S12**: **A** Alluvial plot showing switches to a more open state from 10A to both 10A.TGFB and 10A.shHPA,

**B** Alluvial plot showing switches to a more closed state from 10A to both 10A.TGFB and 10A.shHPA,

**C** Alluvial plot showing switches to a more closed state from 10A to 10A.shHPA and to a more open state from 10A to 10A.TGFB

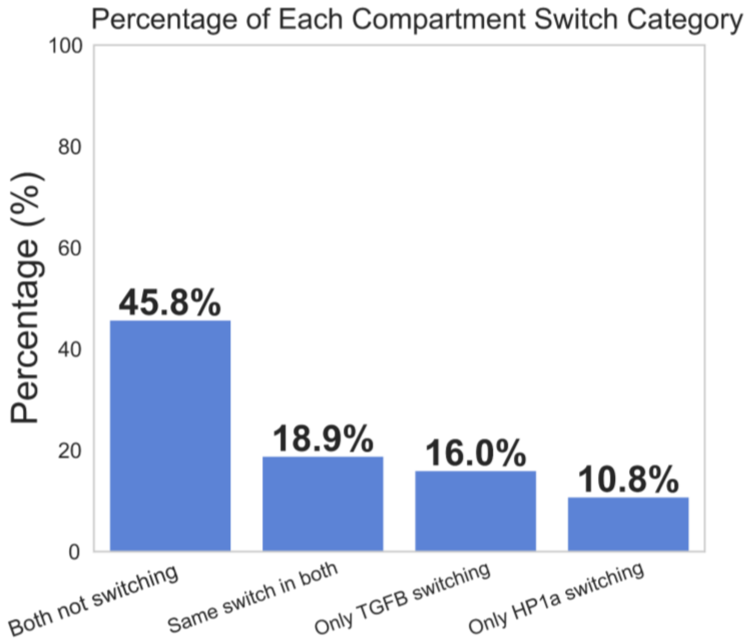

**Figure S13:** Histograms showing subcompartment switches unique to TGFβ-treated cells, HP1α knockdown, and shared between both conditions.

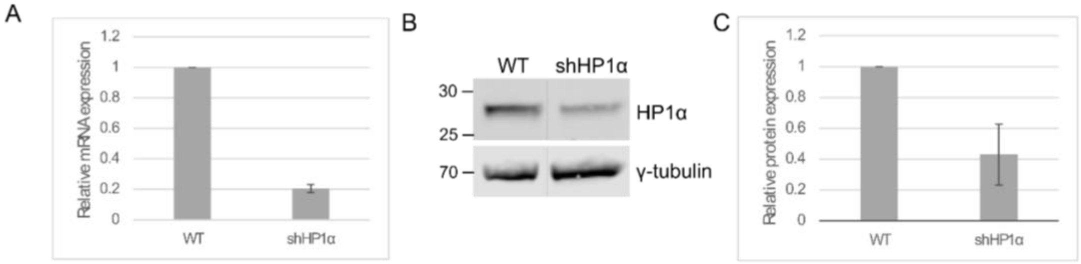

**Figure S14: A** Relative mRNA expression of HP1α in wild-type (WT) and HP1α knockdown (shHP1α) cells.

**B** Representative western blot showing HP1α protein levels in WT and shHP1α cells.γ-tubulin was used as a loading control.

**C** Relative protein expression of HP1α in wild-type (WT) and HP1α knockdown (shHP1α) cells.
